# Luminescence-based complementation assay to assess target engagement and cell permeability of glycolate oxidase (HAO1) inhibitors

**DOI:** 10.1101/2024.06.26.600887

**Authors:** Sabrina R. Mackinnon, Tryfon Zarganes-Tzitzikas, Cassandra J. Adams, Paul E. Brennan, Wyatt W. Yue

## Abstract

Glycolate oxidase (HAO1) catalyses the synthesis of glyoxylate, a common metabolic intermediate that causes renal failure if accumulated. HAO1 inhibition is an emerging treatment for primary hyperoxaluria, a rare disorder of glyoxylate metabolism. Here we report the first cell-based measurement of inhibitor uptake and engagement with HAO1, by adapting the cellular thermal shift assay (CETSA) based on Nano luciferase complementation and luminescence readout. By profiling the interaction between HAO1 and four well-characterised inhibitors in intact and lysed HEK293T cells, we showed that our CETSA method differentiates between low-permeability/high-engagement and high-permeability/low-engagement ligands and is able to rank HAO1 inhibitors in line with both recombinant protein methods and previously reported indirect cellular assays. Our methodology addresses the unmet need for a robust, sensitive, and scalable cellular assay to guide HAO1 inhibitor development and, in broader terms, can be rapidly adapted for other targets to simultaneously monitor compound affinity and cellular permeability.

## 1. Introduction

Inhibition of glycolate oxidase (2-hydroxyacid oxidase, HAO1) has been established as a safe and effective treatment for primary hyperoxaluria type I (PH1)^1^, a rare disorder of glyoxylate metabolism caused by inherited mutations of *AGXT1,* encoding the liver enzyme alanine-glyoxylate aminotransferase, which detoxifies glyoxylate. The primary patho-mechanism of PH1 is toxicity of oxalate, generated from accumulated glyoxylate, which results in deposition of calcium oxalate crystals in renal tissues, leading to end-stage kidney failure^2^. A small interfering RNA targeting HAO1, the enzyme involved in glyoxylate biosynthesis, has recently been approved to treat PH1^3^. Drug discovery campaigns have also been undertaken to identify small molecule HAO1 inhibitors with two lead compounds currently in late stage clinical trials^4,5^.

HAO1 is a flavin mononucleotide (FMN)-dependent enzyme that catalyses the oxidation of glycolate to glyoxylate (EC 1.1.3.15). To date, almost all HAO1 inhibitors target the enzyme’s active site and share a similar physiochemical profile. Specifically, they contain a carboxylic acid moiety to achieve potency by mimicking the enzyme’s substrate glycolate, leading to poor cell permeability and low metabolic stability^6^, and the rest of the molecule is a large hydrophobic moiety interacting with the non-polar residues immediately outside the active site, resulting in poor solubility. Consequently, the success of therapeutic molecules targeting HAO1 is often dependent on a compound’s ability to engage with HAO1 in cells over time, rather than potency. However, cell-based assays for HAO1 inhibitor characterisation published to date measure changes in cell viability or metabolite levels in mouse or CHO cell lines^7–9^, an indirect readout of HAO1 inhibition in non-human cell lines, rather than determining direct engagement of compounds with HAO1 in human cells.

Therefore, an alternative means to quantify compound engagement with HAO1 in cells would be beneficial to future drug discovery efforts. One such type of cell-based target engagement assay – the cellular thermal shift assay (CETSA) – relies on the principle of thermal stabilisation, whereby compound binding decreases the temperature-dependent aggregation of the target protein in a concentration-dependent manner according to its cellular binding affinity^10,11^. The classical CETSA uses Western blot to measure changes to soluble protein levels caused by pre-incubation with compound before heating. Adaptations of the CETSA methodology to increase throughput and flexibility have led to different approaches for quantification of the target protein level, including the SplitLuc complementation system^12,13^.

This reporter-based system relies on the reconstitution of a functional ‘NanoLuc’ luciferase enzyme (engineered from Oluc-19 of the deep sea shrimp *Oplophorus gracilirostris*^14^) by complementation between two separately inactive parts – HiBiT (1.6 kDa), a small 15-amino acid helix attached to the target protein of interest, and LgBiT (17.7 kDa), the remainder of the luciferase protein to be provided in the assay detection reagent along with its substrate furimazine (Figure 1A). Formation of a functional NanoLuc enzyme, generating a luminescent product for assay readout, only takes place when the target protein, and therefore the attached HiBiT helix, is present and able to complement the LgBiT protein provided in the detection reagent to enable turnover of furimazine (Figure 1A). The level of the target protein can then be moderated by temperature (e.g., heat denaturation), and thermal stabilisation of the target protein due to ligand binding can be determined from observed changes in the subsequent luminescence signal^12,13^.

**Figure 1:**
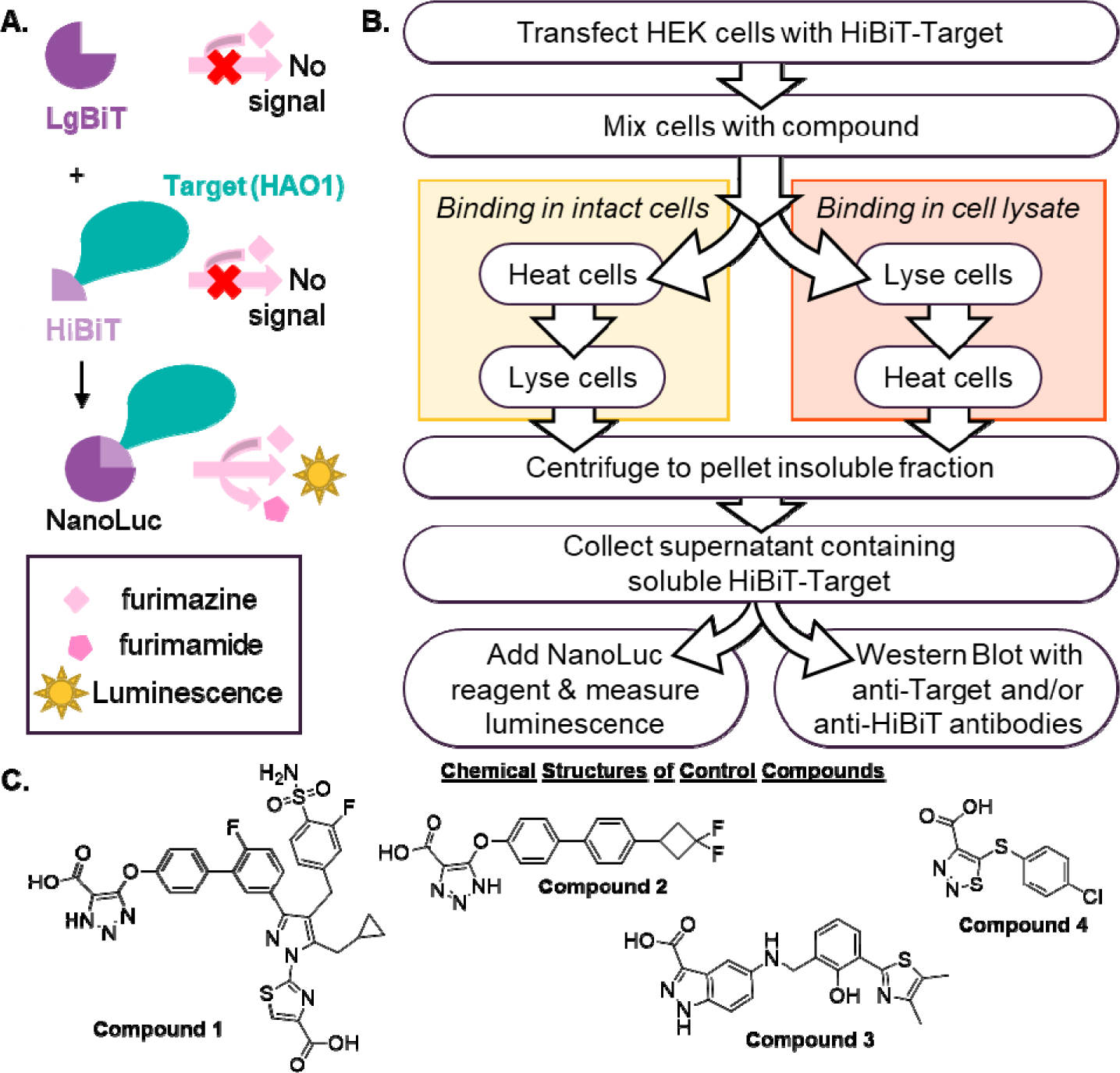
Principle of SplitLuc CETSA. A. Formation of functional NanoLuc luciferase enzyme from LgBiT and HiBiT components to allow turnover of furimazine to furimamide, generating luminescence. B. Schematic of CETSA established in this work. C. Chemical structures of compounds **1**–**4**.

Since its original characterisation^15^ and application to CETSA^12^, HiBiT-LgBiT complementation has been used primarily as a reporter system, where addition of LgBiT is used to locate and/or quantify HiBiT-tagged target protein, in studies monitoring cell processes such as binding and import of viral proteins^16,17^, protein degradation^18,19^ and changes in protein translation^20^. More recently its application in CETSA has been presented^12^, and reported for inhibitor development campaigns^13,21–24^ for a handful of therapeutic targets^25–31^.

To adapt the SplitLuc CETSA for HAO1 drug discovery, we assessed four reported HAO1 inhibitors (Figure 1C) in intact cells and cell lysates. Compound **1** (reported as compound 15 by Ding *et al*.^32^) is a dual glycolate oxidase-lactate dehydrogenase inhibitor that demonstrated high potency and affinity against both recombinant human HAO1 and mouse HAO1 in hepatocytes but had minimal effect in the PH1 mouse model due to poor liver exposure^32^. Compound **2** (reported as compound 36 by Maag *et al*.^4^) has promising *in vitro* and *in vivo* profile^4,33^ and is currently in Phase II/III clinical trials for the treatment of PH1. Compound **3** (reported as compound 29 by Lee *et al*.^34^) is a highly potent HAO1 inhibitor but development of this series was discontinued due to poor cell permeability^34^. Compound **4** (5-[(4-methylphenyl)sulfanyl]-1,2,3-thiadiazole-4-carboxylic acid; CCPST) is a legacy HAO1 inhibitor^35^ found to be unsuitable for therapeutic development due to poor solubility and low cell permeability^1^.

This work describes the adaptation of SplitLuc CETSA to assess cellular permeability and engagement with HAO1 of compounds **1**–**4**. Given the known permeability and cellular stability issues reported for many HAO1 inhibitors, we also investigated whether running CETSA in cell lysates rather than intact cells would allow differentiation between cellular affinity and cellular permeability.

## 2. Material and methods

### 2.1 Recombinant protein production

LgBiT protein, for inclusion in the detection reagent, HiBiT-tagged MBP protein, for use as a positive control, and His-tagged HAO1 protein, for *in vitro* assays and use as a negative control, were expressed and purified from *E. coli* using standard protocols. A description of the plasmids used in this study and details of their expression and purification are provided in the supporting information.

### 2.2 Western Blot of recombinant proteins

Two-fold serial dilutions from 1 mg/mL (20 samples) were prepared for both His-HAO1 and HiBiT-MBP proteins by dilution in 2X SDS-PAGE loading dye. Control samples of either 0.1 mg/mL HiBiT-MBP or 0.1 mg/mL His-HAO1 were also prepared in 2X SDS-PAGE loading dye. Samples were boiled for 5 minutes before loading duplicate 4-12% Bis-Tris SDS-PAGE gels (ThermoFisher) and running gel electrophoresis (1 hour, 180 V) in SDS-MOPS running buffer (ForMedium). Pre-stained molecular weight ladders (Prime-Step™ Prestained Broad Range Protein Ladder; BioLegend) were included in all runs to allow size determination of detected bands. After electrophoresis, one gel was transferred to EZBlue™ Gel Staining Reagent (Sigma Aldrich) and the other was transferred to a PVDF membrane (Bio-Rad Laboratories) for Western Blot using a Trans-Blot Turbo transfer system (Bio-Rad Laboratories) at 25 V for 7 minutes. After blocking with a 5% skim milk solution, membranes were incubated overnight at 4 °C with primary antibodies (anti-HAO1 [1:2000 dilution] or anti-HiBiT [1:2000 dilution]) generated in mouse, before incubation for 4 hours at 4 °C with the secondary antibody, HRP-conjugated anti-mouse IgG (H/L) antibody, generated in goat (antibodies.com; [1:5000 dilution]). Coomassie stained gels and Western Blots treated with SuperSignal West Pico PLUS solution (ThermoFisher) were imaged by white light and chemiluminescence respectively using a BioRad GelDoc XRS+ system.

### 2.3 Chemicals

Compound **4** (4-carboxy-5-[(4-chlorophenyl)sulfanyl]-1,2,3-thiadiazole, CCPST) was purchased from Key Organics (12G-315S). The other three inhibitors were synthesised according to published protocols (compound **1**^32^, compound **2**^4^, compound **3**^34^). Synthetic schemes and detailed synthetic protocols are provided in supplemental information.

### 2.4 *In vitro* characterisation of compounds 1 – 4

Compound binding to HAO1, measured by surface plasmon resonance (SPR); compound inhibition of HAO1, measured by Amplex Red activity assay; and thermal stabilisation of HAO1, measured by differential scanning fluorimetry (DSF), were all determined as described in our previous work^36,37^.

### 2.5 Detection of HAO1 in cells by Western Blot

HEK293T cells (ATCC CRL-3216) were cultured and transfected according to standard protocols, as detailed in the supporting information. Intact cells, 72 hours post-transfection, were diluted to 5 x 10^6^ cells/mL in assay buffer (1XPBS supplemented with 1x Halt protease inhibitors) and lysed by addition of 1% NP-40 (ThermoFisher) followed by rocking at room temperature for 30 minutes. Lysates were centrifuged at 4000 rpm for 10 minutes at 4 °C and an aliquot of each supernatant (soluble protein fraction) sample transferred to an equal volume of SDS-PAGE loading dye and boiled before analysis by SDS-PAGE and Western Blotting as described for recombinant protein samples.

### 2.6 Detection of HAO1 in cells by luminescence

HEK293T cells (ATCC CRL-3216) were cultured and transfected according to standard protocols, as detailed in the supporting information. For temperature range experiments, transfected cells (at 1 x 10^6^ cells/mL) were aliquoted into a 96-well PCR plate (Sarstedt) at 30 µL/well and heated for 3.5 minutes, applying the required temperature gradient across the plate using the VeriFlex technology of a QuantStudio 5 RT-PCR machine (Applied Biosystems), before lysis by addition of 1% NP-40 followed by rocking at room temperature for 30 minutes. For compound experiments requiring intact cells, transfected cells were diluted to 1 x 10^6^ cells/mL in assay buffer and then incubated with varying compound concentrations (12 concentrations, 0–100 μM final concentrations for compounds **1**–**3** and 0– 500 μM final concentrations for compound **4** and 1% final DMSO for all samples) for 1 hour at 37 °C before heating to 55 °C for 3.5 minutes using a QuantStudio 5 RT-PCR machine. After equilibrating to room temperature, cells were lysed by addition of 1% NP-40 and rocking at room temperature for 30 minutes. For compound experiments requiring cell lysates, transfected cells were lysed by addition of 1% NP-40, and incubation with rocking at room temperature for 30 minutes, before 1 hour incubation with compounds at 37 °C and heating to 55 °C for 3.5 minutes.

After lysis, all samples were centrifuged at 4000 rpm for 10 minutes at 4 °C and supernatant (soluble protein fraction) samples transferred to a fresh plate containing an equal volume of CETSA detection reagent – assay buffer supplemented with 100 nM of purified LgBiT-His protein and 0.5X furimazine (Nano-Glo® luciferase assay system, Promega). Samples were then incubated at room temperature for 30 minutes before transfer to a 384-well white plate (Greiner-One) and measurement of luminescence signal using a POLARstar OMEGA plate reader (BMG Labtech). Controls included in all runs were: a standard curve of HiBiT-MBP protein from 0 – 200 nM, recombinant His-HAO1 protein at 1 mg/mL, recombinant LgBiT-His protein at 1 mg/mL and non-transfected HEK293T cells prepared in parallel to the samples being tested. Data were normalized plate-wise using the HiBiT-MBP standard curve. Concentration-response curves were fitted and EC_50_ calculated in GraphPad Prism software using a nonlinear least-squares regression fit to the log (inhibitor) vs response (three parameters) equation.

## 3. Results

### 3.1 SplitLuc reconstitution using recombinant components

The NanoLuc luminescence signal depends upon the complementation of LgBiT protein with a HiBiT-tagged target protein (Figure 1A). We first evaluated NanoLuc luminescence through complementation of a series of control LgBiT proteins and HiBiT-tagged proteins expressed in *E. coli* (Figure S1, Table S1). This includes LgBiT protein incorporated with His_6_-tag at either its N- or C-terminus (His-LgBiT or LgBiT-His), and HiBiT attached to either the N- or C-terminus of maltose binding protein (HiBiT-MBP or MBP-HiBiT). We also included His_6_-tagged HAO1 (His-HAO1, as previously described^36,37^) to ensure HAO1 protein did not interfere with the assay or generate luminescence in the presence of LgBiT and furimazine (Figure S2A).

When measuring the luminescence signal of each protein individually, we observed no signal for His-LgBiT, LgBiT-His or His-HAO1 proteins as expected (Figure S2A). Mixing of one LgBiT protein with either another LgBiT protein or HAO1 protein also does not generate luminescence (Figure S2B), indicating LgBiT alone and LgBiT mixed with HAO1 do not form a functional luciferase, as expected. Unexpectedly, we observed concentration-dependent increase in luminescence with HiBiT-tagged MBP (HiBiT-MBP or MBP-HiBiT) alone (Figure S2A) or when HiBiT-MBP and MBP-HiBiT were mixed together (Figure S2B), prompting us to include LgBiT-free control samples in our optimised assay to allow correction for any background luminescence arising from our HiBiT-tagged HAO1 protein.

As expected, mixing of equimolar amounts of LgBiT and HiBiT proteins together reconstituted functional NanoLuc, generating significant luminescence signal upon addition of the furimazine substrate (Figure S2C). The two HiBiT-tagged proteins (HiBiT-MBP and MBP-HiBiT) lead to similar luminescence profiles when complemented with LgBiT (His-LgBiT or LgBiT-His). By contrast, His-LgBiT and LgBiT-His had distinctly different luminescence profiles when mixed with HiBiT-tagged proteins (Figure S2C). Specifically, His-LgBiT had significantly greater affinity than LgBiT-His for HiBiT-tagged proteins, achieving maximal luminescence by 31.25 nM protein concentration and generating high signal: background ratio at the minimum tested protein concentration (500X greater luminescence at 0.5 nM of each protein compared with 0 nM control). This is in line with previously published parameters, where the concentration of HiBiT positive control proteins was 0.5-1.0 nM^12,25^.

In comparison, our LgBiT-His protein required 5-10 times higher protein concentration to reach this level of luminescence but continued to produce a linear signal up to 125 nM. Based on these results, we chose to include 100 nM of LgBiT-His in our NanoLuc detection reagent, to allow for a wider range of linear luminescence signal. We also confirmed that increasing both HiBiT-tagged MBP concentrations above 100 nM, while keeping LgBiT-His concentration constant, resulted in unchanged luminescence (data not shown).

Since the original CETSA method uses Western blot for quantification of protein levels^38^, we also assessed the suitability of commercially available monoclonal anti-HAO1 (Figure S3) and anti-HiBiT (Figure S4) primary antibodies, towards detection of our recombinant proteins (His-HAO1 and HiBiT-MBP) using a horseradish peroxidase-conjugated secondary antibody and chemiluminescent substrate. The anti-HiBiT antibody did not recognise recombinant His-HAO1 whereas the anti-HAO1 antibody cross-reacted with recombinant HiBiT-MBP at the tested concentrations (1 µg HiBiT-MBP or His-HAO1), and the anti-HiBiT antibody was able to detect much lower protein concentrations (Figures S3-S4).

### 3.2 HiBiT-tagged HAO1, expressed in HEK293T cells, complements recombinant LgBiT

After establishing parameters using recombinant proteins, we measured detection of HiBiT-tagged HAO1 proteins, transiently expressed in HEK293T cells under six different transfection mixes (Table 1; Figure 1B). Based on previous studies reporting significant differences in target protein stability depending on where the HiBiT tag is incorporated^12,13,23^, we compared two different HAO1 constructs, incorporating a HiBiT tag at either the N- or C-terminus (HiBiT-HAO1 or HAO1-HiBiT), which were transfected in the presence or absence of an *AGXT1* plasmid, such that the expressed AGXT1 enzyme would metabolise glyoxylate generated from HAO1 and mitigate any glyoxylate-induced toxicity in cells (transfections II-V, Table 1).

**Table 1:** Plasmid details for tested transfection mixes.

| Transfection number | Plasmid 1 | Plasmid 2 |
| --- | --- | --- |
| I | None | None |
| II | 2.5 µg HiBiT-HAO1 | None |
| III | 1.25 µg HiBiT-HAO1 | 1.25 µg AGXT1-HTStII |
| IV | 2.5 µg HAO1-HiBiT | None |
| V | 1.25 µg HAO1-HiBiT | 1.25 µg AGXT1-HTStII |
| VI | None | 1.25 µg AGXT1-HTStII |

Blank (water instead of DNA) and AGXT1-alone transfections (transfections I and VI respectively) were included as controls. Cells were lysed 72 hours after transfection, and expression of HiBiT-tagged HAO1 was confirmed by Western Blots using anti-HAO1 (Figure 2A) and anti-HiBiT (Figure 2B) primary antibodies. Neither antibody recognised proteins in the control transfections without HiBiT-HAO1 (transfections I and VI), and both Western Blots showed a band of the expected size (42.3 kDa) for at least one replicate of each transfection containing HiBiT-HAO1 (transfections II-V).

**Figure 2:**
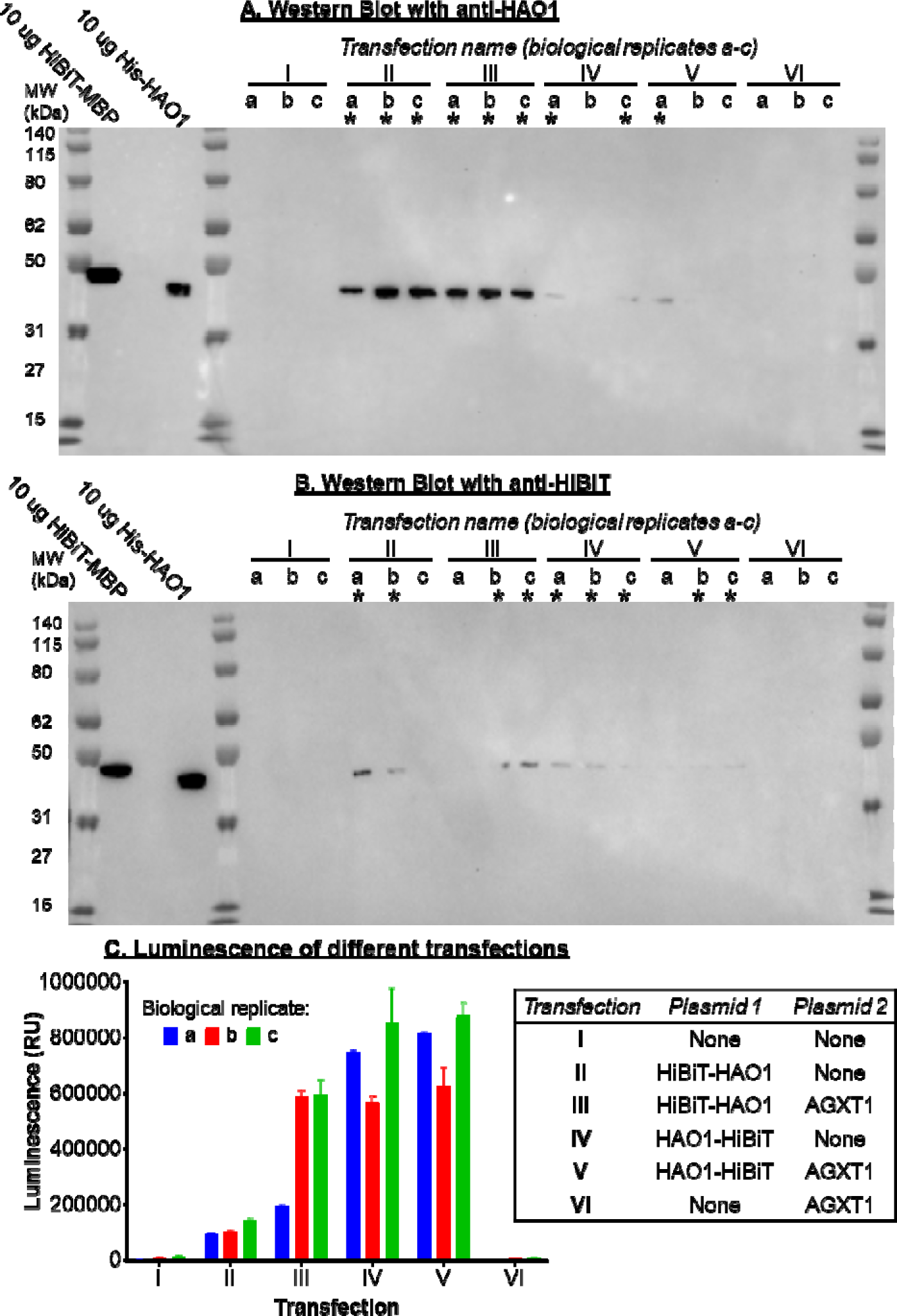
HiBiT-tagged HAO1, expressed in HEK293T, can be detected by Western Blot or by complementation with recombinant LgBiT. Western blot of different transfection combinations, detailed in the inset table, detected with anti-HAO1 (A) or anti-HiBiT monoclonal antibodies (B). Both blots were treated with an HRP-conjugated anti-mouse IgG secondary antibody and imaged after addition of chemiluminescent HRP substrate. Three biological replicates are shown for each transfection mix. C. Luminescence detection of transfections, detailed in inset table, after addition CETSA detection reagent (LgBiT protein and furimazine substrate). Data are represented as mean ± SD for technical triplicates. Results from three biological replicates (a-c) are shown. See also Figures S5 and S8. *Inset: Definition of transfection mixes I–VI*.

Having confirmed expression of HiBiT-tagged HAO1 (transfections II-V) by Western Blot, we tested whether we could detect the HiBiT tag presented by HAO1. We measured luminescence generated upon addition of the CETSA detection reagent (containing LgBiT-His that binds to the HiBiT tag and forms the reconstituted luciferase, and the luciferase substrate furimazine).

As expected, the control transfections lacking a HiBiT tag (transfections I and VI) did not produce a luminescence signal, while the four transfections (transfections II–V) that contained HiBiT-tagged HAO1, all produced luminescence, indicating that NanoLuc complementation had taken place between LgBiT-His protein in the detection reagent and the HiBiT tag presented by HAO1 in cells (Figure 2C, Figure S5). Despite similar or greater protein expression levels detected by Western Blot, transfections containing HiBiT-HAO1 generally produced a lower luminescence signal than transfections containing HAO1-HiBiT, irrespective of the presence of AGXT1 (Figure 2C, Figure S5), indicating that the position of the HiBiT tag relative to HAO1 impacted the resulting luminescence signal. However, further investigation – for example, through co-expression of a fluorescent protein to quantify transfection efficacy allowing direct comparison with the HiBiT-derived signal – is merited to explain the variability seen in these transfections (i.e., replicate a in Figure 2C is of significantly higher luminescence than other transfections with HiBiT-HAO1, despite being treated in an identical manner).

For further experiments, we selected transfection mix V (HAO1-HiBiT + AGXT1, Figure 2C) that contained both glyoxylate metabolising enzymes (HAO1 and AGXT1) while providing the highest luminescence signal.

### 3.3 Decreasing cellular levels of HAO1-HiBiT after heating can be detected by luminescence

To determine the best temperature for monitoring stabilisation induced by compounds, we evaluated the thermal stability of transfected HAO1-HiBiT in HEK293T cells. HEK293T cells, 72 hours post-transfection with HAO1-HiBiT and AGXT1, were heated in either 5 °C increments between 40 and 65 °C or in 2 °C increments between 45 and 65 °C and soluble levels of HAO1-HiBiT were characterised by adding CETSA detection reagent and measuring the luminescence generated (Figure 3). As expected, we observed a temperature-dependent decrease in luminescence, indicative of decreasing levels of soluble HAO1 with increasing temperature. A similar inverse temperature-luminescence relationship was also observed for the other transfections containing HiBiT-tagged HAO1 (Figure S5).

**Figure 3:**
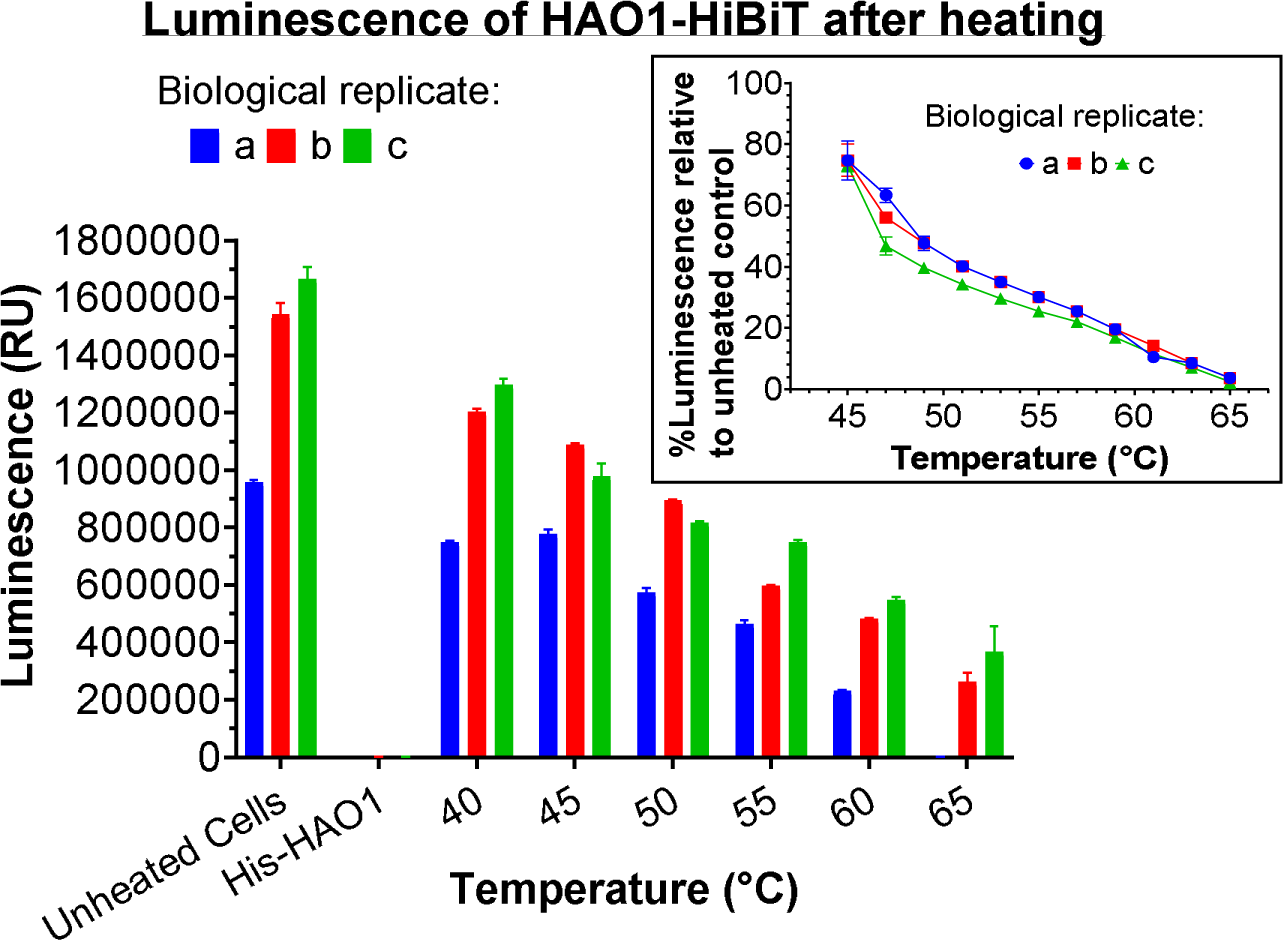
Decreasing cellular levels of HAO1-HiBiT after heating can be detected by luminescence. Detection of soluble levels of HAO1-HiBiT protein, after heating at 2 °C intervals, by luminescence signal after addition of CETSA detection reagent (LgBiT protein and furimazine substrate). Data are represented as mean ± SD for technical triplicates. Results from three biological replicates (a-c) are shown. See also Figure S5.

From these data, we chose 55 °C for all subsequent heating steps as this temperature resulted in a soluble HAO1-HiBiT level that was 35 ± 9% of the unheated control, across six biological replicates, each tested in technical triplicates. We chose this temperature with the aim of unfolding 60-75% of unliganded HAO1 while generating sufficient signal above background that a reduction in stability would also be detected.

### 3.4 Assays of tool compounds with recombinant HAO1 justified CETSA detection of cell-based target engagement

The purpose of the CETSA developed in this work was to evaluate cellular target engagement of four HAO1 inhibitors (compounds **1**-**4**, Figure 4A) for which recombinant enzyme and cell-based activity have been reported (Table S2). We also characterised the compounds for their binding to and inhibition of recombinant HAO1 using our established binding and activity assays^37^, to compare with published profiles (Table S2).

**Figure 4:**
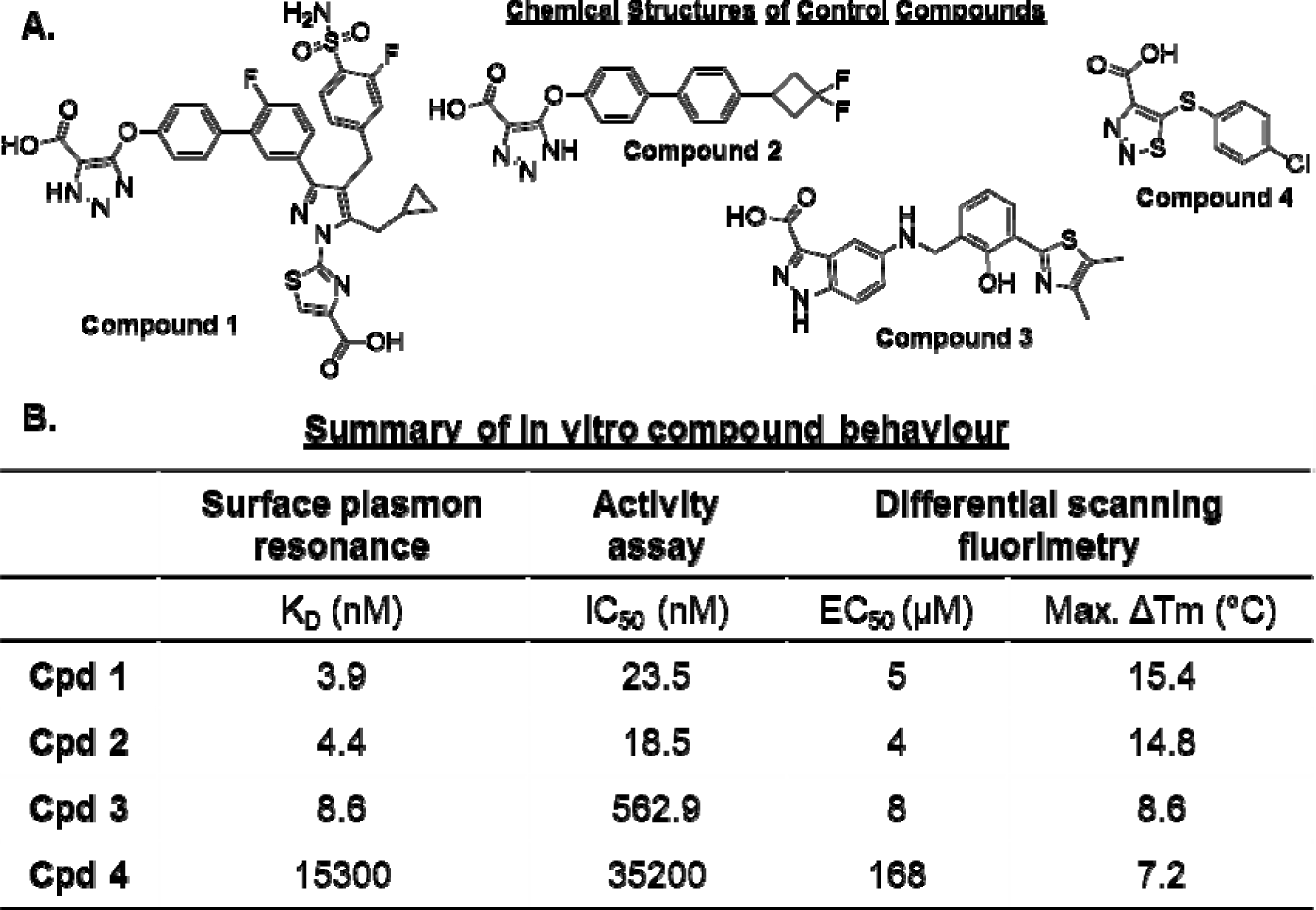
Assays with recombinant HAO1 indicate our chosen compounds are suited to detection of target engagement by CETSA. A. Chemical structures of control compounds. B. Parameters for binding to and inhibition of HAO1 by compounds **1**–**4**, determined in recombinant protein assays. Data are mean values ± SD of three biological replicates. Standard deviations for parameters calculated using a logarithmic scale (IC_50_ and EC_50_) are reported in the text.

Using surface plasmon resonance (SPR), we determined low nanomolar binding affinities (dissociation (binding) constant, *K_D_*) towards HAO1 for compounds **1–3** and a low micromolar binding affinity towards HAO1 for compound **4** (Figure 4, Figure S6A). Binding affinities determined in our SPR assay are in close agreement with published affinities, also determined by SPR, for compound **2** (K_D_ 6.31 nM^33^) and compound **4** (K_D_ 47.5 μM^37^) and represent the first published binding affinities for compounds **1** and **3**.

In our Amplex Red assay monitoring recombinant HAO1 activity, we calculated double-digit nanomolar IC_50_ values for HAO1 inhibition by compounds **1** and **2**, a high nanomolar IC50 value for inhibition by compound **3**, and a low micromolar IC_50_ value for inhibition by compound **4** (pIC_50_ values of 7.6 ± 0.08, 7.7 ± 0.05,6.3 ± 0.08, and4.5 ± 0.06 for compounds **1–4** respectively) (Figure 4, Figure S6B). Our IC_50_ values are, overall, 10-fold less potent than previously reported values (Table S2, also measured in Amplex Red activity assay). The latter discrepancy is commonly observed between independent inhibition measurements due to differences in assay conditions (e.g., protein concentration, reaction time, substrate concentration). Notably, there is clear agreement in relative values between compounds, except that compound **3** appears much less potent in our activity assay than would be expected. This is likely due to interference of the compound with the assay, resulting in oxidation of the Amplex Red dye in the absence of HAO1 activity (data not shown).

Thermal stabilisation of HAO1 upon compound binding is a key pre-requisite for detection of target engagement using CETSA to avoid false negatives, where a ligand does not induce stabilisation^39^ or destabilises the target protein^28^. We previously used differential scanning fluorimetry (DSF) to show that recombinant HAO1 undergoes significant thermal stabilisation upon binding of its cofactor FMN or substrate glycolate^36^. We applied DSF to investigate whether compounds **1**–**4** were able to stabilise recombinant HAO1. Here, HAO1 thermal unfolding was monitored by binding of the hydrophobic fluorescent dye, SYPRO Orange, to exposed regions of the unfolding protein. We observed maximal stabilisation (change in melting temperature, ΔT_m_, at which half of the present protein is unfolded) of recombinant HAO1 by approximately 15 °C for compounds 1 and 2, 9 °C for compound 3 and 7 °C for compound **4**, extrapolating to low micromolar EC_50_ values for compounds **1 – 3** and a mid-micromolar EC50 value for compound **4** (pEC_50_ values of 5.3 ± 0.1, 5.4 ± 0.1, 5.1 ± 0.04, and 3.8 ± 0.1 for compounds **1–4** respectively) (Figure 4, Figure S6C), demonstrating that the degree of thermal stabilisation of HAO1 is proportional to each compound’s affinity and potency values determined from other assays. The observed T_m_ for HAO1 in the absence of added compound was 47 °C, which agrees with the ∼50% reduction in soluble protein observed in transfected HEK293T cells after heating to 49 °C (Figure 3).

Altogether, the reported compound potency and affinity values are reproducible in our assays, indicating a reliable effect on HAO1, and we demonstrated that thermal stabilisation can be used to evaluate and rank HAO1 inhibitors in terms of *in vitro* affinity. This also provides confidence in the concept of increased HAO1 thermal stability upon inhibitor binding in cells.

### 3.5 Our CETSA approach reliably measured HAO1 inhibitor affinity and permeability

We began exploring compound-induced stabilisation of HAO1 in HEK293T by performing the six transfection combinations as described above and comparing soluble HAO1 levels after heating to 55 °C, with and without preincubation with 200 µM compound **2** (chosen for its high affinity and relative ease of synthesis). We observed a significant increase (ranging from 60 ± 2% to 183 ± 8%) in soluble levels of HAO1 protein for all transfections including either HiBiT-HAO1 or HAO1-HiBiT (transfections II–V), and no change (±3%) in the low background luminescence for control transfections without HiBiT-HAO1/HAO1-HiBiT (transfections I and VI) under the same conditions (Figure 5A).

**Figure 5:**
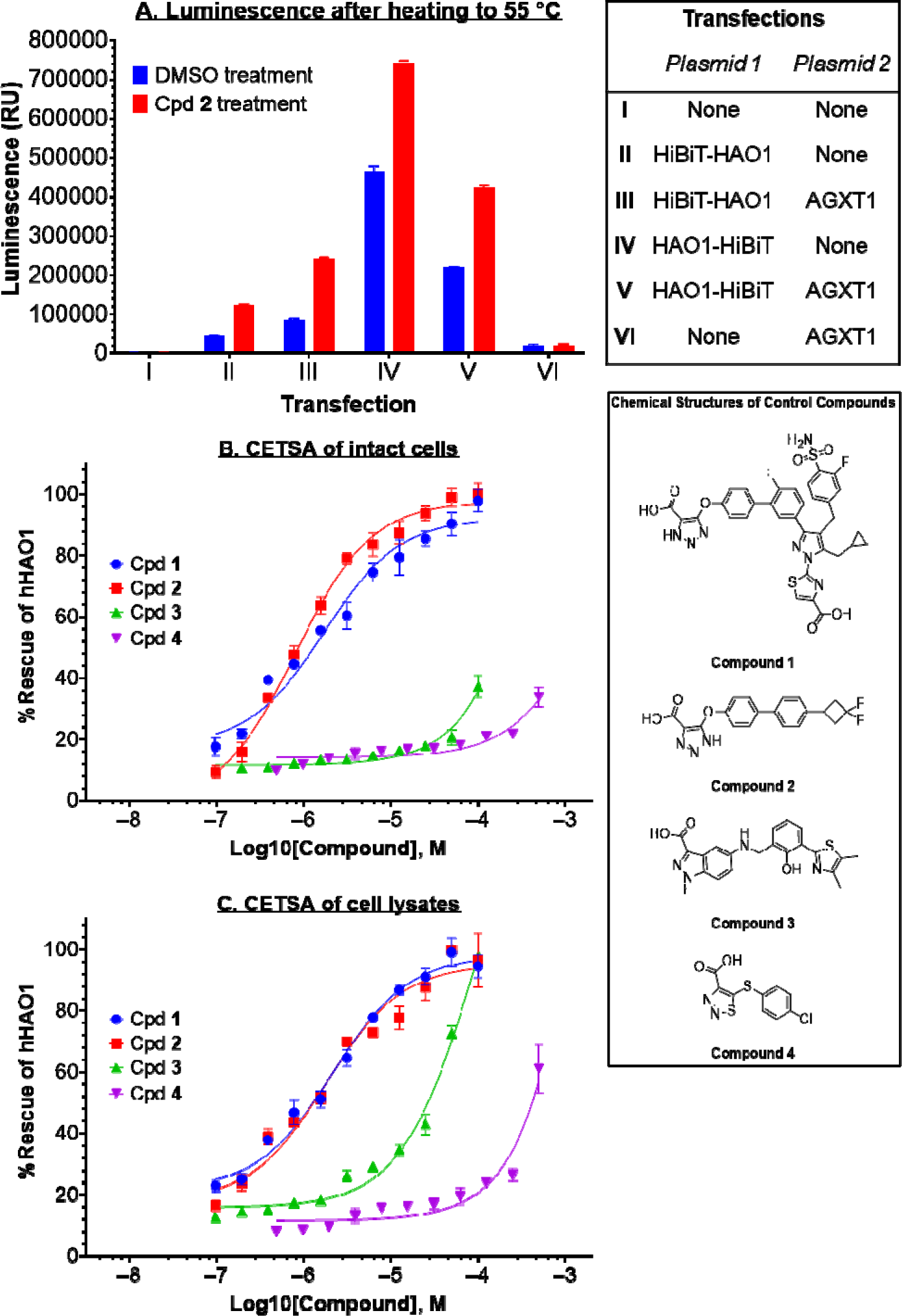
Our CETSA assay differentiates between poor affinity and poor permeability of our chosen compounds. Data are represented as mean ±SD of technical triplicates. A. Luminescence detection of transfections, detailed in inset table (top), after preincubation with 200 μM compound **2** or equivalent DMSO and heating to 55 °C (intact cells). *Inset (top): Definition of transfection mixes I*–*VI*. B – C. Luminescence detection of soluble HAO1-HiBiT after preincubation with increasing compound concentrations and heating to 55 °C, measured in either intact cells (B) or cell lysates (C). *Inset (middle): Chemical structures of control compounds*. D. Summary of compound parameters determined in CETSA experiments. See also Figure S7.

We next determined soluble HAO1 levels after heating to 55 °C, following preincubation with different concentrations of each of the four compounds. We titrated a range of compound concentrations (0–100 μM for compounds **1**–**3** and 0–500 μM for compound **4**) to HEK293T cells co-transfected with HAO1-HiBiT and AGXT1 (72 hours prior to addition of compounds) and incubated at 37 °C for an hour, then heating samples to 55 °C and quantifying soluble HAO1 protein levels by luminescence. We observed a maximal concentration of soluble HAO1, relative to an unheated control sample, of 98 ± 4%, 100 ± 4%, 37 ± 4% and 34 ± 3% for cells treated with compounds **1**–**4** respectively (Figure 5B). This means that there was no unfolding of HAO1 protein after heating when cells were treated with high concentrations (> 25 μM) of compound **1** or **2**, while > 50% of HAO1 was unfolded after heating when cells were treated with the maximum tested concentrations of compound **3** and **4** (i.e., 100 μM compound **3** or 500 μM compound **4**). The calculated EC_50_ values towards HAO1 stabilisation were 1.7 μM (pEC_50_ 5.8 ± 0.13), 0.9 μM (pEC_50_ 6.1 ± 0.05), >100 μM and >500 μM for compounds **1**–**4** respectively (Figure 5D, biological replicate in Figure S7A).

While our EC_50_s for compounds **1** and **2** are approximately 20-fold weaker than their published cellular IC_50_s (calculated from measurement of oxalate reduction in mouse hepatocytes; Table S2), the two compounds demonstrate similar behaviour in our CETSA, as observed in cell-based assays measuring reduction of oxalate in hyperoxaluric mouse hepatocytes^32,33^. The observed differences in apparent efficacy between the two cell-based assays can be attributed to two key factors. Firstly, temperature has a significant effect on both protein integrity and compound binding affinity and our CETSA involves heating of samples to 55 °C after pre-incubation at 37 °C, while samples in the reported mouse hepatocyte assays are kept constant at 37 °C. Secondly, our assay directly measures how much compound has bound to, and stabilised, HAO1, whereas the hepatocyte-based assay measured a phenotypic outcome – oxalate reduction – that is dependent not just on HAO1 inhibition but on the compound’s overall effect on oxalate production in the cell, including changes to the activity of other enzymes in the glyoxylate metabolic pathway (e.g., lactate dehydrogenase, which converts glyoxylate to oxalate). Previous studies comparing compound parameters determined by CETSA and other cell-based assays have also showed that the determined EC_50_ values are often an order of magnitude weaker in CETSA, but that the ranking of compounds is consistent across assays^31,38^. Interestingly, our CETSA EC_50_ values for compounds **1** and **2** were essentially the same as those obtained in DSF – most likely because DSF was performed at high recombinant protein concentrations (4.7 µM His-HAO1), approximately two orders of magnitude higher than the expression level in CETSA (<50 nM HAO1-HiBiT at 55 °C).

In contrast, we observed very weak interaction between HAO1 and compounds **3** and **4** in our CETSA (Figure 5B), indicating stabilisation of HAO1 in cells was at least two orders of magnitude weaker than observed in our DSF assay (Figure 4B, Figure S6C), and does not reflect the low- to mid-micromolar cellular efficacy reported for these compounds, particularly the nanomolar potency of compound **3** in recombinant protein assays (Table S2).

Given the solubility and permeability issues previously reported for compounds **3**^34^ and **4**^1^, we investigated whether cell lysis prior to heating, thus bypassing the need for compounds to be cell-permeable, would reconcile this apparent discrepancy between the weak target engagement observed in CETSA and the higher affinity/potency observed in other cell-based assays and recombinant protein assays. We reasoned that cell lysis would have minimal impact on the observed engagement of compounds that were able to reach HAO1 in intact cells – such as compounds **1** and **2** – while causing a significant improvement in the observed engagement of compounds prevented from reaching HAO1 in cells due to poor permeability.

We determined EC_50_ values of the 4 compounds as described above, except that cells were lysed prior to preincubation with compound. When cells were lysed prior to compound treatment, we observed a maximal concentration of soluble HAO1, relative to an unheated control sample, of 95 ± 4%, 97 ± 9%, 98 ± 1% and 61 ± 8% (Figure 5C), resulting in EC_50_ values of 2.2 μM (pEC_50_ 5.7 ± 0.07), 1.8 μM (pEC_50_ 5.7 ± 0.14), 104 μM (pEC_50_ 4.0 ± 0.14) and ∼400 μM (unable to calculate pEC_50_) for cells treated with compounds **1**–**4** respectively (Figure 5D; biological replicate in Figure S7B). Therefore, in line with our hypothesis, compounds **1** and **2** maintained their maximal stabilisation and efficacy in cell lysates compared to intact cells, while compounds **3** and **4** showed significant improvement of maximal stabilisation reached and calculated EC_50_ values, in cell lysates compared to intact cells.

## 4. Discussion

We optimised and validated the SplitLuc CETSA approach to generate a robust, sensitive, and scalable method for early-stage drug discovery. This enables us to characterise the engagement between HAO1 and inhibitors in human cells by monitoring protein thermal stabilisation, and to deconvolute cellular affinity from cellular permeability of HAO1 inhibitors by comparing their engagement with HAO1 in intact cells and cell lysates. This work represents the first report characterising HAO1 inhibitors in a human cell line and, furthermore, is the first assay directly quantifying engagement of HAO1 with compound, rather than measuring the downstream changes in metabolite levels resulting from HAO1 inhibition, in any cell line.

We first synthesised four reported HAO1 inhibitors, compounds **1**–**4**, and characterised their inhibition of and binding to recombinant HAO1 in established biophysical assays. Compounds **1**, **2** and **3** demonstrated low nanomolar affinity for HAO1, while compound **4** is of double-digit micromolar affinity, as previously reported. We next compared thermal stability, in the presence and absence of these compounds, for recombinant HAO1 measured by DSF, and for HAO1 present in HEK293T intact cells or cell lysate measured by CETSA. We observed strong correlation in melting temperature (T_m_; at which half the present HAO1 protein is unfolded) between DSF (47 °C) and CETSA (49 °C), indicating that the thermal stability of recombinant HAO1 is a good indicator of HAO1 stability in human cells.

In line with this observation, the calculated EC_50_s for thermal stabilisation of HAO1 by compounds **1** and **2** are nearly identical when measured by DSF, intact cell CETSA and cell lysate CETSA. We attributed the similar efficacy values to the high protein concentration in DSF relative to CETSA, which balances out possible factors leading to lower efficacy in the latter assay (e.g., cellular crowding, off-target interactions, compound metabolism).

Compounds **3** and **4**, however, show discrepancies between DSF and CETSA calculated EC_50_s. For compound **4**, the calculated EC_50_s for thermal stabilisation of HAO1 is similar when measured by either DSF or cell lysate CETSA (168 and 400 µM respectively) but is significantly weaker when measured by intact cell CETSA, such that an EC_50_ could not be reliably determined. This indicates that compound **4** cannot effectively cross the cell membrane to interact with HAO1, consistent with previous reports of low permeability and limited aqueous solubility of this compound^1^.

In line with our observations of compounds **1** and **2**, we recorded single-digit nanomolar affinity by SPR and a single-digit micromolar EC_50_ by DSF for compound **3**. However, similar to compound **4**, the observed thermal stabilisation in intact cell CETSA was significantly weaker than in DSF which prevented EC_50_ calculation, in agreement with the reported suboptimal permeability of this compound^34^. Surprisingly, the thermal stabilisation of HAO1 by compound **3** was not completely rescued by lysing the cells prior to addition of compounds and heating, indicating that poor permeability alone did not account for the weaker cellular affinity of compound **3**. Additional factors that may account for differences between binding to recombinant HAO1 and binding to HAO1 in a cellular environment include binding to serum protein, precipitation of compound within the cell due to poor solubility, and metabolic degradation of the compound before reaching the target protein – all of which are known risks associated with carboxylic acid containing structures.

While our CETSA assay provides a better indication of compound cellular efficacy than previously reported indirect assay in non-human cell lines, its current application in HEK293T cells could introduce non-physiological levels of HAO1 expression and differences in the cellular environment compared to hepatocytes, where therapeutic molecules in development need to engage with HAO1. Future work adapting SplitLuc CETSA for use in the immortalised hepatoma-derived cell line HepG2, which has been recently validated as a model cell line for primary hyperoxaluria^40^, would overcome this limitation. An additional avenue to explore in future would be the relationship between compound binding to HAO1 (measured in the developed CETSA), compound inhibition of HAO1 (measured in a cellular environment) and compound phenotypic effect (i.e., reduction of oxalate), including the identification of any indirect mechanisms reducing oxalate (e.g., inhibition of oxalate synthesis by LDHA, upregulation of glyoxylate detoxification by AGXT1).

In addition to providing an improved cellular assay for the characterisation of HAO1 inhibitors, this work also elaborates on the general use of the LgBiT-HiBiT complementation system by generating two LgBiT protein reagents, one with high sensitivity but a narrow linear range (His-LgBiT), and another with lower sensitivity but a broad linear range (LgBiT-His). Additionally, we generated recombinant HiBiT-tagged MBP proteins to eliminate the need to purchase expensive positive control peptide or protein for assay validation. Furthermore, our observation of decreasing luminescence at high protein concentrations has not been previously reported for the HiBiT-LgBiT system and is an important consideration in deciding upon the concentration of LgBiT protein in the detection reagent and of any HiBiT-tagged control samples. Lastly, most guidance on CETSA to date suggests trialling HiBiT-tag at either terminus of the target protein to identify the best signal when complementing with LgBiT. Here, we elaborated on this consideration, noting some variations in the detection of HiBiT-tagged target protein that depend on the site of HiBiT attachment. We reason that, unlike in previously reported examples^12,13,23^, these signal differences are due to occlusion of the HiBiT tag, rather than changes in thermal stability of the target protein. Due consideration should therefore be made on how HiBiT accessibility could be influenced by homo- or hetero-oligomerisation of the target protein (e.g. HAO1 is a homotetramer that can form octamers in solution, Figure S8).

## 5. Conclusions and Outlook

We have developed a robust, scalable, and direct cellular assay, the first reported in a human cell line, to facilitate HAO1 inhibitor development. Our adapted SplitLuc CETSA reliably ranked four reported HAO1 inhibitors and differentiated among compounds with high affinity and good permeability (e.g., compounds **1** and **2**), compounds with poor permeability (e.g., compound **4**), and compounds with high affinity but poor cellular efficacy due to more complex factors (e.g., compound **3**). We also show that comparison of HAO1 thermal stabilisation in DSF, intact cell CETSA and cell lysate CETSA can differentiate between low affinity, poor permeability and more complex causes of limited cellular efficacy, providing a cost-effective and scalable approach to guide HAO1 drug discovery. Given the body of work demonstrating the easy adaptation of CETSA to a broad range of targets, our methodology can similarly be applied to a broad range of drug discovery projects to characterise target engagement, cellular permeability and overall cellular efficacy of lead compounds in a single assay readout. We view this would be particularly helpful during the hit-to-lead optimisation stage of novel drug discovery, to inform the decision making process for whether or not to advance promising compounds towards determination of ADME and PK properties in animal models.

## Supporting Information

Methods, figures and tables relating to preparation and validation of tool proteins. Detailed experimental procedures for compound synthesis, cell culture and in vitro assays. Supplemental figures showing biological replicates of data reported in main figures.

## Acknowledgements

Plasmid for HAO1 expression in E. coli was generously provided by Structural Genomics Consortium University of Oxford. This work was supported by a Harrington Principal Investigator Award (to W.W.Y.) issued by Harrington Discovery Institute at University Hospitals in Cleveland, Ohio, USA, through its UK charity, Fund For Cures UK, Ltd. We thank the Wellcome Trust for supporting the earlier stages of this work (092809/Z/10/Z and 106169/ZZ14/Z). C.J.A. gratefully acknowledges funding from the NIHR Oxford Biomedical Research Centre (BRC), University of Oxford. T.Z.T and P.E.B. thank Alzheimer’s Research UK for support (ARUK-2021DDI-OX).

## Author contributions

Conceptualization, W.W.Y and S.R.M; Methodology, S.R.M and T.Z; Investigation, S.R.M and T.Z; Visualisation, S.R.M. and T.Z; Writing – Original Draft, W.W.Y and S.R.M; Writing – Review & Editing, W.W.Y, S.R.M, T.Z and C.J.A; Funding Acquisition - W.W.Y, S.R.M, T.Z and C.J.A; Supervision, W.W.Y and P.E.B.

