## Supplemental data for "Luminescence-based complementation assay to assess target engagement and cell permeability of glycolate oxidase (HAO1) inhibitors"

1. Supplementary Methods.......................................................................................................2
   1.1. Plasmid details
   1.2. Recombinant protein production
   1.3. Culture and transfection of HEK293T cells
   1.4. Synthesis of compounds **1** – **3**
   1.5. *In vitro* characterisation of compounds **1** – **4**
2. Supplementary Figures and Tables.....................................................................................21
   - Figure S1. Purification of recombinant protein SplitLuc components and controls.
   - Figure S2. Recombinant HiBiT-tagged proteins, but not His-HAO1, complement His-tagged LgBiT.
   - Figure S3 Detection of recombinant His-HAO1, separated using SDS-PAGE, by Coomassie staining and Western Blot.
   - Figure S4 Detection of recombinant HiBiT-MBP, separated using SDS-PAGE, by Coomassie staining and Western Blot.
   - Figure S5 Luminescence detection of HiBiT-tagged HAO1 from different transfections after heating between 40 and 65 °C.
   - Figure S6: In vitro inhibition, binding and stabilisation of recombinant His-HAO1 by control compounds.
   - Figure S7: Biological replicates of concentration-response testing of compounds 1–4 in SplitLuc CETSA.
   - Figure S8. Structural models of possible oligomers of HiBiT-tagged HAO1 and the spinach homologue of HAO1, sGOX.
   - Table S1: Vector and sequence details of constructs used in this study.
   - Table S2: Published characterisation data for compounds **1** – **4** in recombinant protein and cellular assays.
3. Supplemental References....................................................................................................32
1. **Supplemental Methods**
   1.1. **Plasmid details***.*

DNA synthesis of full length HAO1 (residues M1–I370) constructs with a HiBiT tag (GSVSGWRLFKKISGS) incorporated at either the N-terminus or the C-terminus in an ampicillin resistant mammalian expression vector (pHTBV1.1), generating HiBiT-HAO1 and HAO1-HiBiT constructs respectively, and of *E. coli* expression constructs (HiBiT-MBP, MBP-HiBiT, and LgBiT-His in pNIC28-CTH0 vector and His-LgBiT in pNIC28-Bsa4 vector), was performed by Twist Bioscience. The *E. coli* expression construct for His-tagged near full-length HAO1 (residues M1-S368; in pNIC28-Bsa4) was kindly provided by SGC, University of Oxford. A full length AGXT1 (residues M1–L392) construct was generated by subcloning a gene string of full length *AGXT1* (synthesised by GenScript) into an ampicillin resistant mammalian expression vector containing a C-terminal sequence to add a TEV protease cleavage site, twin-Strep affinity tag and His6 tag (pHTBV1.1-CT10H-SIII-LIC vector, available upon request from SGC, University of Oxford). This was achieved by performing inverse PCR to linearise the chosen vector and PCR amplification of the AGXT1 gene string to produce products with complementary overhangs, followed by assembly of the two products using the In-Fusion system (Clontech) and verification of the assembled plasmid’s sequence by Sanger sequencing. Amino acid sequences of the constructs used in this study are provided in Table S1. All *E. coli* expression plasmids conferred kanamycin resistance whereas all mammalian expression plasmids conferred ampicillin resistance.

1.2. **Recombinant protein production.**

Each *E. coli* expression construct was transformed into *E. coli* BL21(DE3) cells (ThermoFisher) according to the manufacturer’s instructions and incubated overnight at 37 °C, with shaking at 200 rpm, in Lysogeny Broth media (ForMedium) supplemented with 50 μg/mL kanamycin and 34 μg/mL chloramphenicol. For each construct, 5 mL of overnight culture was added to 1 L of auto-induction Terrific Broth media (ForMedium), prepared according to the manufacturer’s instructions, and supplemented with 50 μg/mL kanamycin and 34 μg/mL chloramphenicol. Cells were incubated for 24 hours at 30 °C before harvesting by centrifugation (4000 x g, 15 minutes, 4 °C). Cell pellets were resuspended in base buffer (50 mM HEPES, pH 7.5, 500 mM NaCl, 5% glycerol, 0.5 mM TCEP), lysed by sonication (programmed to run 5 seconds on, 10 seconds off, 35% amplitude, for 15 minutes) and then centrifuged to remove insoluble material (18000 x g, 1 hour, 4 °C). For purification of HAO1 protein, 0.1 mM FMN was added prior to lysis. Affinity purification was performed by addition of 1-3 mL of Ni-NTA resin (ThermoFisher), pre-equilibrated in base buffer, to each supernatant and subsequent rotation at 4 °C for 1 hour, followed by batch purification with sequential washes of base buffer supplemented with 20 mM and 40 mM imidazole before elution of His-tagged proteins in base buffer supplemented with 250 mM imidazole. Fractions containing the protein of interest were identified by SDS-PAGE analysis and pooled for further purification using size exclusion chromatography, performed in base buffer using a Superdex 75 Increase column (GE Healthcare) attached to a BioRad NGC chromatography system (BioRad Laboratories). For SDS-PAGE analysis, 10 µL of sample was added to 5 µL 2X SDS-PAGE loading dye (4% SDS, 0.25 M Tris, pH 6.8, 20% glycerol, 10% β-mercaptoethanol, and 0.01% Bromophenol Blue) and boiled for 5 minutes before loading onto a 4-12% Bis-Tris SDS-PAGE gel (ThermoFisher) and running gel electrophoresis (1 hour, 180 V) in SDS-MOPS buffer (ForMedium). Final protein pools, made up of fractions containing the protein of interest, were identified by combined examination of the resultant chromatograms and SDS-PAGE analysis (See Figure S1). Purified proteins were aliquoted, snap frozen in liquid nitrogen and stored at – 80 °C.

1.3. **Culture and transfection of HEK293T cells.**

HEK293T (human embryonic kidney) cells (female line) were obtained from ATCC (CRL-3216) and maintained in DMEM media (Gibco) supplemented with 10% foetal bovine serum and 100 U/mL penicillin/streptomycin (Invitrogen), in incubators maintained at 37 °C with 8% CO2 and 95% humidity. Prior to transfection, 5x10^5^ cells were aliquoted per well into 6-well dishes and returned to the incubator overnight before transfection by washing the cells with PBS, and addition of 1.2 mL Opti-MEM media (ThermoFisher) to each well and then 125 µL of the appropriate transfection complex – consisting of 1.875 µL Lipofectamine 3000, 5 µL P3000 reagent, and 2.5 µg DNA prepared in Opti-MEM media – was added dropwise to each well. Where two plasmids were transfected, 1.25 µg of each was included in the transfection mix. After 3 hours further incubation, Opti-MEM and transfection solutions were replaced with 2.5 mL D-MEM media and incubated for 72 hours prior to experimental procedures. Three biological replicates of each transfection were performed in parallel, and each plate included a non-transfected control.

1.4. **Synthesis of compounds 1 – 3.**

**1.4.1. General synthetic methods***.*

All commercially obtained solvents and reagents were used as received. All solvents used for chemical reactions were anhydrous grade, unless specifically indicated. Structures of the target compounds in this work were assigned by use of NMR and MS spectroscopy. ^1^H NMR spectra were recorded on a Bruker Advance III (400MHz) or a Varian 400MR (400MHz) NMR spectrometer. Chemical shifts are reported in parts per million (ppm, d) using residual solvent line as an internal reference. Splitting patterns are reported as s (singlet), d (doublet), t (triplet), q (quartet), m (multiplet), or br s (broad singlet). Coupling constants (J) are reported in hertz (Hz). Unless specifically indicated, chromatography refers to a flash chromatography on a silica gel column.

**1.4.2. Chemistry synthetic schemes.**

**Scheme 1**: Synthesis of Compound **1**

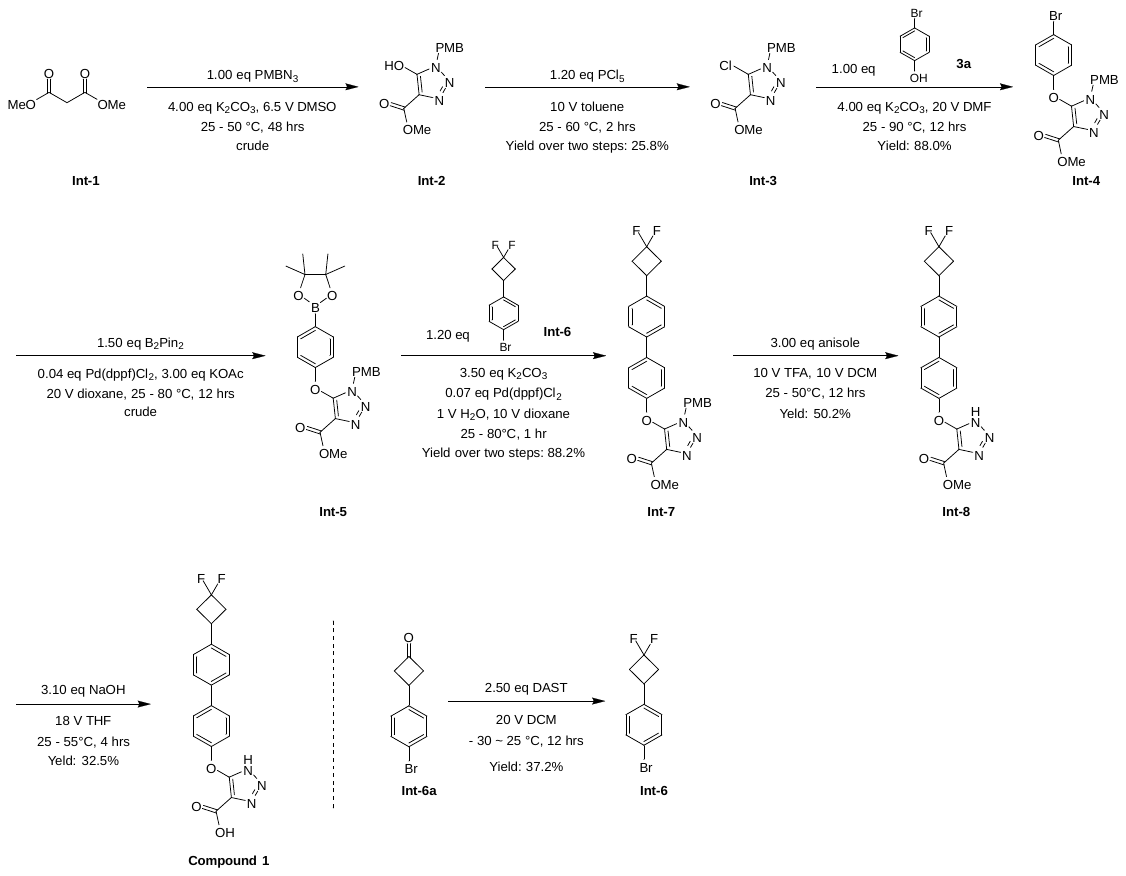

**Scheme 2**: Synthesis of Compound **2**

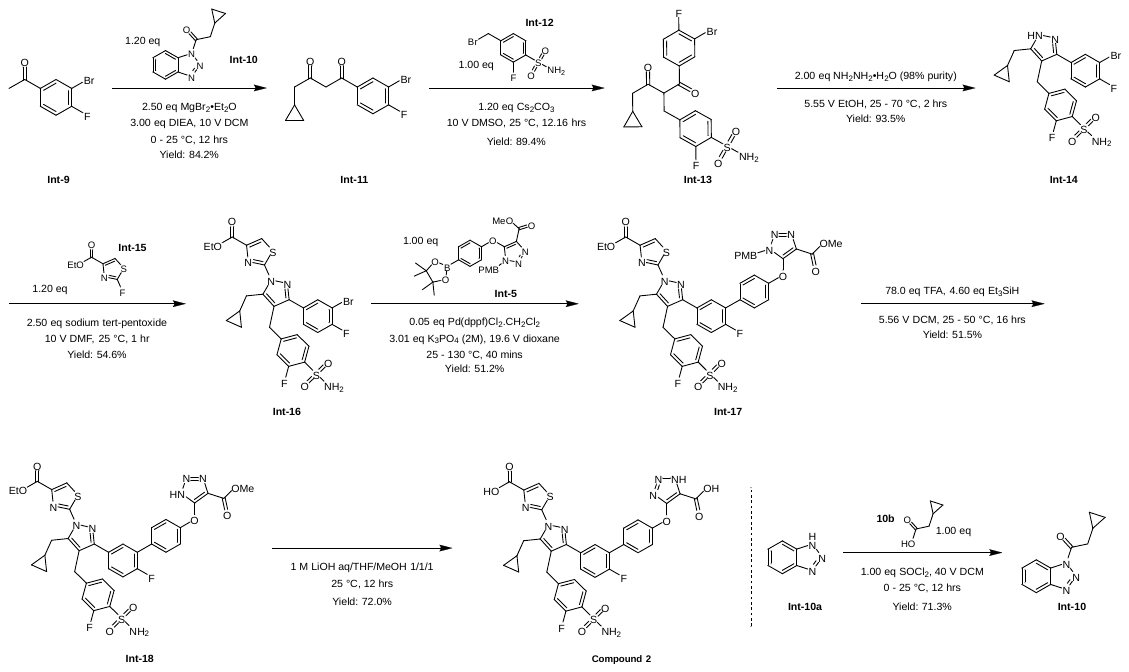

**Scheme 3**: Synthesis of Compound **3**

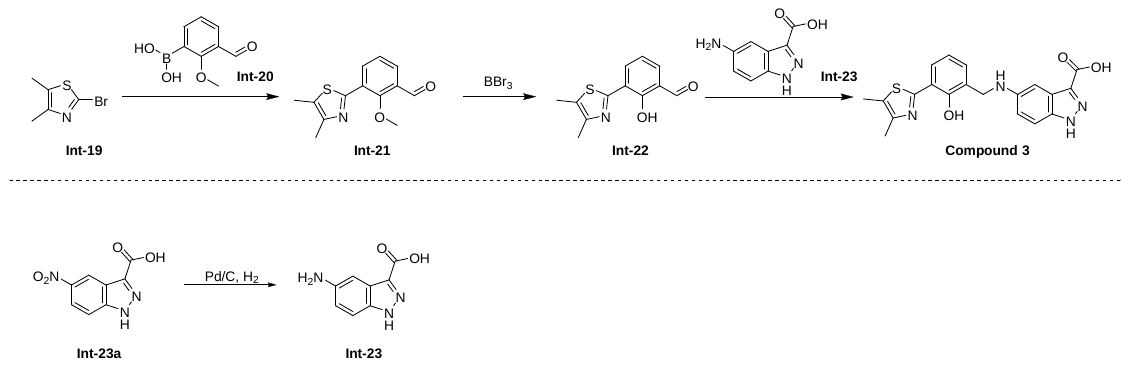

**1.4.3 Synthesis of Key Intermediates**

**Synthesis of methyl 5-hydroxy-1-(4-methoxybenzyl)-1H-1,2,3-triazole-4-carboxylate (Intermediate 2).** To a solution of PMBN_3_ (9.00 g, 55.2 mmol, 1.00 *eq*) in DMSO (58.5 mL) was added **Int-1** (9.98 g, 75.6 mmol, 8.68 mL, 1.37 *eq*) and K_2_CO_3_ (30.5 g, 221 mmol, 4.00 *eq*) at 25 °C, the mixture was heated to and stirred at 50 °C for 48 hours. TLC (Petroleum ether : Ethyl acetate = 2: 1) showed PMBN_3_ (R_f_ = 0.70) was consumed, and one main spot (R_f_ = 0.30) was formed. The mixture was diluted with water (150 mL) and then washed with methyl tert-butyl ether (100 mL * 2). The organic layers were discarded, the aqueous phase was adjusted to pH 3 with 3 M HCl below 10 °C, and then it was extracted with ethyl acetate (100 mL * 3). Combined organic layers were washed with brine (100 mL * 3), dried with Na_2_SO_4_, filtered and concentrated. **Int-2** (12.2 g, crude) was obtained as yellow oil. The crude product was used to next step without purification. **^1^H NMR**: (400 MHz, DMSO-*d_6_* *δ* 7.20 (d, *J* = 8.4 Hz, 2H), 6.92 - 6.89 (m, 2H), 5.25 (s, 2H), 3.76 (s, 3H), 3.73 (s, 3H).

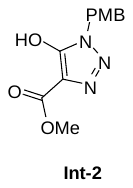

**Synthesis of methyl 5-chloro-1-(4-methoxybenzyl)-1H-1,2,3-triazole-4-carboxylate (Intermediate 3).** To a solution of **Int-2** (10.2 g, 38.8 mmol, 1.00 *eq*) in toluene (102 mL) was added PCl_5_ (9.68 g, 46.5 mmol, 1.20 *eq*) at 25 °C, the mixture was heated to and stirred at 60 °C for 12 hours. TLC (Petroleum ether : Ethyl acetate = 2 : 1) showed some of **Int-2** (R_f_ = 0.31) remained and desired spot (R_f_ = 0.25) was formed. The mixture was cooled to 25 °C, poured into ice NH_3_.H_2_O (100 mL), and then it was extracted with ethyl acetate (100 mL * 3). Combined organic layers were washed with brine (100 mL), dried with Na_2_SO_4_, filtered and concentrated. The residue was purified by column chromatography (SiO_2_, Petroleum ether : Ethyl acetate = 100 : 0 to 1 : 1, Petroleum ether : Ethyl acetate = 2 : 1, R_f_ = 0.25). **Int-3** (2.81 g, 9.98 mmol, 25.8% yield) was obtained as a yellow solid. **^1^H NMR**: (400 MHz, CDCl_3_) *δ* 7.30 - 7.27 (m, 2H), 6.92 - 6.89 (m, 2H), 5.51 (s, 2H), 3.97 (s, 3H), 3.80 (s, 3H).

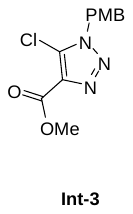

**Synthesis of methyl 5-(4-bromophenoxy)-1-(4-methoxybenzyl)-1H-1,2,3-triazole-4-carboxylate (Intermediate 4).** To a solution of **Int-3** (2.80 g, 9.94 mmol, 1.00 *eq*) in DMF (46.0 mL) was added K_2_CO_3_ (5.50 g, 39.8 mmol, 4.00 *eq*) and **Int-3a** (1.72 g, 9.94 mmol, 1.00 *eq*) at 25 °C, and the reaction mixture was heated to and stirred at 90 °C for 12 hours. TLC (Petroleum ether : Ethyl acetate = 2 : 1) indicated **Int-3** (R_f_ = 0.25) was consumed completely and a main spot (R_f_ = 0.40) was formed. The reaction mixture was cooled to 25 °C, then poured into water (150 mL) and extracted with ethyl acetate (150 mL * 2). Combined organic layers were washed with brine (100 mL * 3), dried over Na_2_SO_4_, filtered and concentrated. The residue was purified by column chromatography (SiO_2_, Petroleum ether : Ethyl acetate = 100 : 1 to 25 : 1, Petroleum ether : Ethyl acetate = 2 : 1, R_f_ = 0.40). **Int-4** (3.66 g, 8.75 mmol, 88.0% yield) was obtained as light yellow solid. **^1^H NMR**: (400 MHz, DMSO-*d_6_*) *δ* 7.50 - 7.47 (m, 2H), 7.17 (d, *J* = 8.8 Hz, 2H), 6.89 - 6.83 (m, 4H), 5.44 (s, 2H), 3.71 (s, 3H), 3.61 (s, 3H).

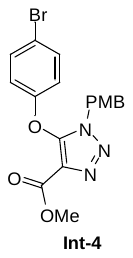

**Synthesis of methyl 1-(4-methoxybenzyl)-5-(4-(4,4,5,5-tetramethyl-1,3,2-dioxaborolan-2-yl)phenoxy)-1H-1,2,3-triazole-4-carboxylate (Intermediate 5).** To a solution of **Int-4** (3.36 g, 8.03 mmol, 1.00 *eq*) in dioxane (68.0 mL) was added B_2_Pin_2_ (3.06 g, 12.1 mmol, 1.50 *eq*), KOAc (2.37 g, 24.1 mmol, 3.00 *eq*), Pd(dppf)Cl_2_.CH_2_Cl_2_ (262 mg, 321 μmol, 0.04 *eq*) at 25 °C. The mixture was degassed and purged with N_2_ 3 times, then it was heated to and stirred at 80 °C for 12 hours under N_2_. LC-MS showed **Int-4** was consumed and the desired MS (Rt = 0.667 min) was detected. The reaction mixture was cooled to 25 °C, then poured into water (60.0 mL) and extracted with ethyl acetate (60.0 mL * 2). Combined organic layers were washed with brine (50.0 mL * 1), dried over Na_2_SO_4_, filtered and concentrated. The residue was purified by column chromatography (SiO_2_, Petroleum ether : Ethyl acetate = 100 : 1 to 5 : 1, Petroleum ether : Ethyl acetate = 2 : 1, R_f_ = 0.40). **Int-5** (3.74 g, crude) was obtained as light yellow oil. **LC-MS**: product: Rt = 0.667 min, m/z = 446.1 (M+H)^+^. **^1^H NMR**: (400 MHz, DMSO-*d_6_*) *δ* 7.61 (d, *J* = 8.4 Hz, 2H), 7.16 (d, *J* = 8.4 Hz, 2H), 6.86 (dd, *J* = 19.2, 8.8 Hz, 4H), 5.44 - 5.39 (m, 2H), 3.69 (s, 3H), 3.59 (s, 3H), 1.28 (s, 12H).

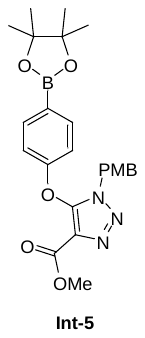

**Synthesis of 1-bromo-4-(3,3-difluorocyclobutyl)benzene (Intermediate 6).** To a solution of **Int-6a** (1.50 g, 6.66 mmol, 1.00 *eq*) in DCM (30.0 mL) was added DAST (1.07 g, 6.66 mmol, 880 μL, 1.00 *eq*) at -30 °C. The mixture was warmed to and stirred at 25 °C for 12 hours. TLC (Petroleum ether : Ethyl acetate = 10 : 1) indicated **Int-6a** (R_f_ = 0.40) remained and a main spot (R_f_ = 0.68) was formed. The mixture was poured into 10.0% NaHCO_3_ solution (50.0 mL) at 0 °C, then it was separated, and the aqueous phase was extracted with DCM (50.0 mL * 2). Combined organic layers was dried with Na_2_SO_4_, filtered and concentrated. The residue was purified by column chromatography (SiO_2_, Petroleum ether : Ethyl acetate = 100 : 0 to 50 : 1, Petroleum ether : Ethyl acetate = 10 : 1, R_f_ = 0.68). **Int-6** (612 mg, 2.48 mmol, 37.2% yield) was obtained as colourless oil. **^1^H NMR**: (400 MHz, CDCl_3_ ) *δ* 7.48 - 7.46 (m, 2H), 7.12 (d, *J* = 8.4 Hz, 2H), 3.39 - 3.31 (m, 1H), 3.05 - 2.98 (m, 2H), 2.71 - 2.61 (m, 2H). **^19^F NMR**: (400 MHz, CDCl_3_) *δ* - 82.44 ~ - 81.93, - 99.43 ~ - 98.91.

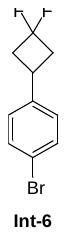

**Synthesis of methyl 5-((4'-(3,3-difluorocyclobutyl)-[1,1'-biphenyl]-4-yl)oxy)-1-(4-methoxybenzyl)-1H-1,2,3-triazole-4-carboxylate (Intermediate 7).** To a solution of **Int-5** (1.11 g, 2.39 mmol, 1.00 *eq*) in dioxane (11.0 mL) and H_2_O (1.00 mL) was added **Int-6** (707 mg, 2.86 mmol, 1.20 *eq*), K_2_CO_3_ (1.15 g, 8.35 mmol, 3.50 *eq*), Pd(dppf)Cl_2_.CH_2_Cl_2_ (136 mg, 167 μmol, 0.07 *eq*) at 25 °C. The mixture was degassed and purged with N_2_ 3 times, and then the mixture was heated to and stirred at 80 °C under N_2_ for 1 hour. LC-MS showed **Int-5** was consumed and the desired MS (Rt = 0.676 min) was detected. The reaction mixture was cooled to 25 °C and concentrated. The residue was purified by column chromatography (SiO_2_, Petroleum ether : Ethyl acetate = 100 : 0 to 4 : 1, Petroleum ether : Ethyl acetate = 2 : 1, R_f_ = 0.20). **Int-7** (1.06 g, 2.10 mmol, 88.2% yield) was obtained as a brown solid. **LC-MS**: product: Rt = 0.676 min, m/z = 506.2 (M+H)^+^. **^1^H NMR**: (400 MHz, DMSO-*d_6_*) *δ*7.70 - 7.59 (m, 4H), 7.49 - 7.38 (m, 2H), 7.19 (d, *J* = 8.8 Hz, 2H), 7.01 - 6.97 (m, 2H), 6.87 - 6.84 (m, 2H), 5.44 (s, 2H), 3.72 - 3.69 (m, 3H), 3.62(s, 3H), 3.50 - 3.4(m, 1H), 3.08 - 2.97(m, 2H), 3.77 - 2.64(m, 2H).

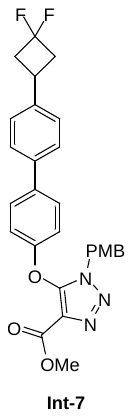

**Synthesis of methyl 5-((4'-(3,3-difluorocyclobutyl)-[1,1'-biphenyl]-4-yl)oxy)-1H-1,2,3-triazole-4-carboxylate (Intermediate 8).** To a solution of **Int-7** (500 mg, 824 μmol, 1.00 *eq*) in DCM (4.00 mL) was added anisole (268 mg, 2.47 mmol, 269 μL, 3.00 *eq*) and TFA (7.68 g, 67.3 mmol, 5.00 mL, 81.6 *eq*) at 25 °C. The mixture was heated to and stirred at 50 °C for 12 hours. LC-MS showed **Int-7** was consumed and the desired MS (Rt = 0.624 min) was detected. The reaction mixture was cooled to 25 °C and concentrated. The residue was purified by prep-TLC (SiO_2_, Petroleum ether : Ethyl acetate = 1 : 1, R_f_ = 0.19). **Int-8** (220 mg, 414 μmol, 50.2% yield, 72.5% purity) was obtained as a light yellow solid. **LC-MS**: product: Rt = 0.624 min, m/z = 386.0 (M+H)^+^.

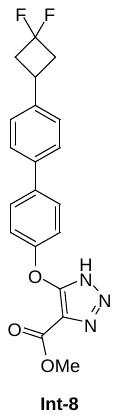

**Synthesis of 5-((4'-(3,3-difluorocyclobutyl)-[1,1'-biphenyl]-4-yl)oxy)-1H-1,2,3-triazole-4-carboxylic acid (Compound 1).** To a solution of **Int-8** (220 mg, 414 μmol, 1.00 *eq*) in THF (4.00 mL) was added a solution of 1M NaOH (51.3 mg, 1.28 mmol, 3.10 *eq*) in water. The mixture was stirred at 55 °C for 4 hours. LC-MS showed **Int-8** was consumed, and the desired MS (Rt = 0.894 min) was detected. The reaction mixture was cooled to 25 °C, then poured into water (10.0 mL) and extracted with methyl tert-butyl ether (10.0 mL * 3), the aqueous phase was adjusted to pH 4 with 1 M HCl, then it was extracted with ethyl acetate (10.0 mL * 3). Combined organic layers were dried over Na_2_SO_4_, filtered and concentrated. The residue was purified by prep-HPLC (column: Phenomenex luna C18 150*25mm*10µm;mobile phase: [water(FA)-ACN];B%: 41%-71%,10 minutes), and the eluent was concentrated at 45 °C and lyophilized. **Compound 1** (50.0 mg, 134 μmol, 32.5% yield, 99.8% purity) was obtained as a white solid. **LC-MS**: product: Rt = 0.894 min, m/z = 372.1 (M+H)^+^. **HPLC**: product: Rt = 3.363 mins, 99.8% purity under 220 nm. **^1^H NMR**: (400 MHz, DMSO-*d_6_*) *δ* 7.66 - 7.61 (m, 4H), 7.39 (d, *J* = 8.4 Hz, 2H), 7.15 - 7.13 (m, 2H), 3.43 (br s, 1H), 3.06 - 3.00 (m, 2H), 2.71 - 2.67 (m, 2H). **^19^F NMR**: (400 MHz, DMSO-*d_6_*) *δ* - 80.79 ~ - 80.28, - 97.31 ~ 96.81.

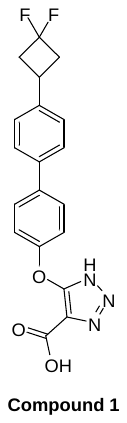

**Synthesis of 1-(1H-benzo[d][1,2,3]triazol-1-yl)-2-cyclopropylethan-1-one (Intermediate 10).** To a solution of **Int-10a** (23.8 g, 199 mmol, 4.00 *eq*) in DCM (200 mL) was added SOCl_2_ (5.94 g, 49.9 mmol, 3.62 mL, 1.00 *eq*) at 25 °C, and the mixture was stirred at 25 °C for 0.5 hours. **Int-10b** (5.00 g, 49.9 mmol, 1.00 *eq*) was added dropwise to the mixture at 0 °C, and the mixture was warmed to and stirred at 25 °C for 11.5 hours. TLC (Petroleum ether : Ethyl acetate = 2 : 1) showed **Int-10b** (R_f_ = 0.55) was consumed and one new spot (R_f_ = 0.72) was formed. The mixture was filtered, the filter cake was washed with DCM (50.0 mL * 3), the filtrate was washed with saturated NaHCO_3_ (100 mL) and separated. Organic layer was washed with saturated NaHCO_3_ (50.0 mL), brine (50.0 mL), dried with Na_2_SO_4_, filtered and concentrated. The residue was purified by column chromatography (SiO_2_, Petroleum ether : Ethyl acetate = 100 : 0 to 50 : 1, Petroleum ether : Ethyl acetate = 5 : 1, R_f_ = 0.70). **Int-10** (7.17 g, 35.6 mmol, 71.3% yield) was obtained as colourless oil. **^1^H NMR**: (400 MHz, DMSO-*d_6_*) *δ* 8.26 - 8.23 (m, 2H), 7.80 - 7.86 (m, 1H), 7.63 - 7.60 (m, 1H), 3.34 - 3.32 (s, 2H), 1.25 - 1.22 (m, 1H), 0.59 - 0.57 (m, 2H), 0.35 - 0.32 (m, 2H).

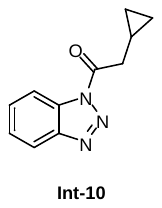

**Synthesis of 1-(3-bromo-4-fluorophenyl)-4-cyclopropylbutane-1,3-dione (Intermediate 11).** To a solution of **Int-10** (6.79 g, 33.7 mmol, 1.20 *eq*), and **Int-9** (6.10 g, 28.1 mmol, 1.00 *eq*) in DCM (61.0 mL) was added MgBr_2_.Et_2_O (18.1 g, 70.3 mmol, 2.50 *eq*) in one portion. The mixture was cooled to 0 °C and DIEA (10.9 g, 84.3 mmol, 14.7 mL, 3.00 *eq*) was added dropwise, then the mixture was warmed to and stirred at 25 °C for 12 hours. TLC (Petroleum ether : Ethyl acetate = 10 : 1) showed most of **Int-9** (R_f_ = 0.49) was consumed and one main spot (R_f_ = 0.58) was formed. The reaction was cooled to 0 °C, then 3 M HCl was added to the mixture dropwise until pH was 3. The reaction mixture was then extracted with DCM (50.0 mL * 2), and the organic layers were washed with brine (100 mL), dried with Na_2_SO_4_, filtered and the filtrate was concentrated. The residue was purified by column chromatography (SiO_2_, Petroleum ether : Ethyl acetate = 1 : 0, Petroleum ether : Ethyl acetate = 10 : 1, R_f_ = 0.58). **Int-11** (7.08 g, 23.7 mmol, 84.2% yield) was obtained as yellow oil. **^1^H NMR**: (400 MHz, DMSO-*d_6_*) *δ* 8.29 (dd, *J* = 6.8, 2.0 Hz, 1H), 8.06 - 8.02 (m, 1H), 7.55 (t, *J* = 8.4 Hz, 1H), 3.36 (s, 2H), 2.34 (d, *J* = 7.2 Hz, 2H), 1.08 - 1.05 (m, 1H), 0.53 - 0.50 (m, 2H), 0.22 - 0.20 (m, 2H).

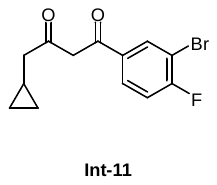

**Synthesis of 4-(2-(3-bromo-4-fluorobenzoyl)-4-cyclopropyl-3-oxobutyl)-2-fluoro benzenesulfonamide (Intermediate 13).** To a solution of **Int-11** (1.30 g, 4.35 mmol, 1.00 *eq*) in DMSO (13.0 mL) was added Cs_2_CO_3_ (1.70 g, 5.22 mmol, 1.20 *eq*) at 25 °C, and the mixture was stirred at 25 °C for 10 minutes before addition of **Int-12** (1.17 g, 4.35 mmol, 1.00 *eq*) to the mixture in portions, and the mixture was stirred at 25 °C for 12 hours. TLC (Petroleum ether : Ethyl acetate = 1 : 1) indicated **Int-12** (R_f_ = 0.42) was consumed completely and a main spot (R_f_ = 0.52) was formed. The reaction mixture was diluted with ethyl acetate (20.0 mL) and filtered to remove the solids. The filtrate was adjusted to pH 4 with 1 M HCl, and diluted with water (30.0 mL), then extracted with ethyl acetate (20.0 mL * 2). Combined organic layer was washed with saturated NH_4_Cl (30.0 mL * 3). The combined organic layer was dried with Na_2_SO_4_, filtered and concentrated. The residue was purified by column chromatography (SiO_2_, Petroleum ether : Ethyl acetate = 100 : 1 to 5 : 1, Petroleum ether : Ethyl acetate = 2 : 1, R_f_ = 0.34). **Int-13** (1.89 g, 3.89 mmol, 89.4% yield) was obtained as yellow oil. **^1^H NMR**: (400 MHz, DMSO-*d_6_*) *δ* 8.31 (dd, *J* = 6.8, 2.4 Hz, 1H), 8.07 - 8.03 (m, 1H), 7.65 (t, *J* = 8.0 Hz, 1H), 7.57 (s, 2H), 7.55 - 7.53 (m, 1H), 7.38 (d, *J* = 11.6 Hz, 1H), 7.24 (d, *J* = 8.0 Hz, 1H), 5.43 (t, *J* = 7.2 Hz, 1H), 3.32 - 3.18 (m, 2H), 2.46 - 2.43 (m, 2H), 0.88 - 0.83 (m, 1H), 0.40 - 0.38 (m, 2H), -0.01 - -0.03 (m, 2H).

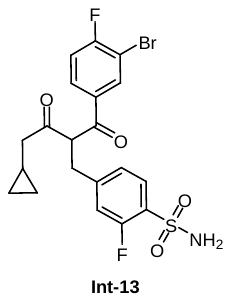

**Synthesis of 4-((3-(3-bromo-4-fluorophenyl)-5-(cyclopropylmethyl)-1H-pyrazol-4-yl)methyl)-2-fluorobenzenesulfonamide (Intermediate 14).** To a solution of **Int-13** (360 mg, 740 μmol, 1.00 *eq*) in ethanol (2.00 mL) was added NH_2_NH_2_.H_2_O (75.6 mg, 1.48 mmol, 73.3 μL, 98.0% purity, 2.00 *eq*) at 25 °C, and the mixture was heated to and stirred at 70 °C for 2 hours. LC-MS showed **Int-13** was consumed and the desired MS (Rt = 0.607 min) was detected. The mixture was cooled to 25 °C and concentrated under vacuum. The crude product was used to next step without purification. **Int-14** (334 mg, 692 μmol, 93.5% yield) was obtained as a light yellow foam. **LC-MS**: product: Rt = 0.607 min, m/z = 483.9 (M+H)^+^. **^1^H NMR**: (400 MHz, CDCl_3_) *δ* 7.81 (t, *J* = 8.0 Hz, 1H), 7.65 (dd, *J* = 6.4, 2.0 Hz, 1H), 7.32 - 7.28 (m, 1H), 7.10 (d, *J* = 8.0 Hz, 1H), 7.03 (br d, *J* = 8.0 Hz, 1H), 6.91 (br d, *J* = 10.8 Hz, 1H), 3.99 (s, 2H), 2.48 (d, *J* = 7.2 Hz, 2H), 0.99 - 0.95 (m, 1H), 0.64 - 0.60 (m, 2H), 0.24 - 0.21 (m, 2H). **^19^F NMR**: (400 MHz, CDCl_3_) *δ* - 108.0, - 110.9.

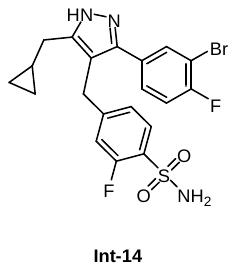

**Synthesis of ethyl 2-(3-(3-bromo-4-fluorophenyl)-5-(cyclopropylmethyl)-4-(3-fluoro-4-sulfamoylbenzyl)-1H-pyrazol-1-yl)thiazole-4-carboxylate (Intermediate 16).** To a solution of **Int-14** (330 mg, 684 μmol, 1.00 *eq*) and **Int-15** (144 mg, 821 μmol, 1.20 *eq*) in DMF (3.00 mL) was added sodium 2-methylbutan-2-olate (188 mg, 1.71 mmol, 2.50 *eq*) at 25 °C, and the mixture was stirred at 25 °C for 1 hour. LC-MS showed **Int-14** was consumed and the desired MS (Rt = 0.701 min) was detected. The mixture was diluted with ethyl acetate (10.0 mL), then it was washed with water (10.0 mL * 3). The organic layer was washed with brine (10.0 mL), dried with Na_2_SO_4_, filtered and concentrated. The residue was purified by prep-TLC (SiO_2_, Petroleum ether : Ethyl acetate = 1**:**1, R_f_ = 0.36). **Int-16** (245 mg, 374 μmol, 54.6% yield, 97.2% purity) was obtained as a light yellow foam. **LC-MS**: product: Rt = 0.701 min, m/z = 637.0 (M+H)^+^.

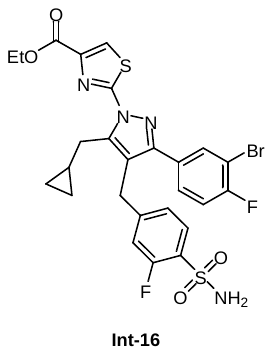

**Synthesis of ethyl 2-(5-(cyclopropylmethyl)-3-(6-fluoro-4'-((1-(4-methoxybenzyl)-4-(methoxycarbonyl)-1H-1,2,3-triazol-5-yl)oxy)-[1,1'-biphenyl]-3-yl)-4-(3-fluoro-4-sulfamoylbenzyl)-1H-pyrazol-1-yl)thiazole-4-carboxylate (Intermediate 17).** To a solution of **Int-16** (253 mg, 386 μmol, 1.00 *eq*) and **Int-5** (179 mg, 386 μmol, 1.00 *eq*) in dioxane (5.00 mL) was added K_3_PO_4_ (2 M, 0.580 mL, 3.01 *eq*) solution and Pd(dppf) Cl_2_.CH_2_Cl_2_ (15.8 mg, 19.3 μmol, 0.050 *eq*) at 25 °C, and the mixture was stirred at 25 °C for 10 minutes under N_2_. The mixture was heated to and stirred at 130 °C for 30 minutes under N_2_. LC-MS showed 6.58% of **Int-16** (Rt = 0.646 min) remained and the desired MS (Rt = 0.709 min) was detected. The mixture was concentrated. The residue was purified by prep-HPLC (column: Waters Xbridge 150*25 mm*5µm;mobile phase: [water (ammonia hydroxide v/v) -ACN];B%: 61%-91%, 9 min), and peak 2 was concentrated. **Int-17** (180 mg, 197 μmol, 51.2% yield, 98.2% purity) was obtained as a light yellow solid. **LC-MS**: product: Rt = 0.709 min, m/z = 896.3 (M+H)^+^. **^1^H NMR**: EW39702-12-P2 (400 MHz, DMSO-*d_6_*) *δ* 8.37 (s, 1H), 7.67 - 7.55 (m, 5H), 7.38 - 7.36 (m, 3H), 7.21 - 7.16 (m, 3H), 7.08 (br d, *J* = 8.8 Hz, 1H), 7.01 (d, *J* = 8.8 Hz, 2H), 6.85 (d, *J* = 8.8 Hz, 2H), 5.45 (s, 2H), 4.35 - 4.30 (m, 2H), 4.20 - 4.16 (m, 2H), 3.67 (s, 3H), 3.63 (s, 3H), 3.18 (br d, *J* = 6.8 Hz, 2H), 1.33 (t, *J* = 6.8 Hz, 3H), 1.16 - 1.13 (m, 1H), 0.35 (br d, *J* = 7.2 Hz, 2H), 0.26 (br d, *J* = 4.4 Hz, 2H).

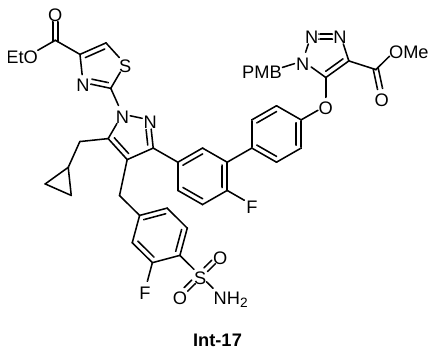

**Synthesis of ethyl 2-(5-(cyclopropylmethyl)-3-(6-fluoro-4'-((4-(methoxycarbonyl)-1H-1,2,3-triazol-5-yl)oxy)-[1,1'-biphenyl]-3-yl)-4-(3-fluoro-4-sulfamoylbenzyl)-1H-pyrazol-1-yl)thiazole-4-carboxylate (Intermediate 18).** To a solution of **Int-17** (180 mg, 201 μmol, 1.00 *eq*) in DCM (1.00 mL) was added Et_3_SiH (72.4 mg, 623 μmol, 99.5 μL, 3.10 *eq*) at 25 °C. The mixture was stirred for 2 minutes at 25 °C. TFA (1.21 g, 10.7 mmol, 791 μL, 53.0 *eq*) was added to the mixture. The reaction was heated to and stirred at 50 °C for 4 hours. LC-MS showed 64.1% of **Int-17** (Rt=0.701 min) remained and trace desired MS (Rt = 0.640 min) was detected. The mixture was cooled to 25 °C, Et_3_SiH (35.0 mg, 301 μmol, 48.1 μL, 1.50 *eq*) and TFA (573 mg, 5.02 mmol, 373 μL, 25.0 *eq*) were added to the mixture at 25 °C, then it was heated to and stirred at 50 °C for 12 hours. LC-MS showed **Int-17** was consumed and the desired MS (Rt = 0.668 min) was detected. The reaction mixture was cooled to 25 °C, then it was concentrated. The residue was purified by prep-TLC (SiO_2_, Petroleum ether : Ethyl acetate = 1 : 1, R_f_ = 0.15). **Int-18** (87.0 mg, 104 μmol, 51.5% yield, 92.3% purity) was obtained as a light yellow solid. **LC-MS**: product: Rt = 0.668 min, m/z = 776.3 (M+H)^+^.

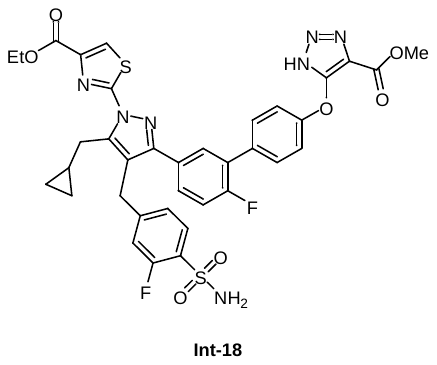

**Synthesis of 2-(3-(4'-((5-carboxy-1H-1,2,3-triazol-4-yl)oxy)-6-fluoro-[1,1'-biphenyl]-3-yl)-5-(cyclopropylmethyl)-4-(3-fluoro-4-sulfamoylbenzyl)-1H-pyrazol-1-yl)thiazole-4-carboxylic acid (Compound 2).** To a solution of **Int-18** (87.0 mg, 104 μmol, 1.00 *eq*) was added a solution of LiOH (22.2 mg, 925 μmol, 8.94 *eq*) in H_2_O (1.00 mL), THF (1.00 mL) and MeOH (1.00 mL), and the mixture was stirred at 25 °C for 12 hours. LC-MS showed **Int-18** was consumed and the desired MS (Rt = 0.879 min) was detected. The reaction mixture was cooled to 25 °C and concentrated. The residue was purified by prep-HPLC (column: Phenomenex Luna C18 150*25 mm*10µm; mobile phase: [water (FA) -ACN];B%: 39%-69%, 10 min), and the eluent was concentrated at 45 °C and lyophilized. **Compound 2** (55.0 mg, 74.5 μmol, 72.0% yield, 99.4% purity) was obtained as an off-white solid. **LC-MS**: product: Rt = 0.897 min, m/z = 734.0 (M+H)^+^. **HPLC**: product: Rt = 1.291 mins, 99.4% purity under 220 nm. **^1^H NMR**: (400 MHz, DMSO-*d_6_*) *δ* 8.31 (s, 1H), 7.67 (t, *J* = 8.4 Hz, 1H), 7.62 - 7.59 (m, 4H), 7.42 (s, 1H), 7.40 - 7.34 (m, 2H), 7.18 (br d, *J* = 12 Hz, 1H), 7.14 (br d, *J* = 8.8 Hz, 2H), 7.08 (d, *J* = 8.4 Hz, 1H), 4.20 (s, 2H), 3.18 (br d, *J* = 7.2 Hz, 2H), 1.17 - 1.12 (m, 1H), 0.38 - 0.33 (m, 2H), 0.25 - 0.23 (m, 2H). **^19^F NMR**: (400 MHz, DMSO-*d_6_*) *δ* - 111.1, - 117.9.

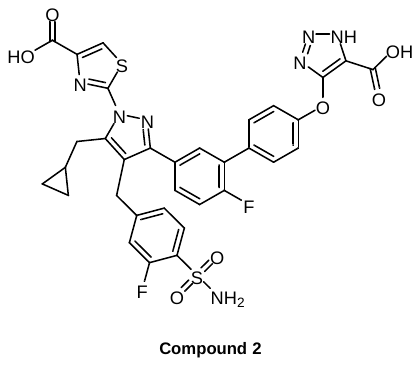

**Synthesis of 3-(4,5-dimethylthiazol-2-yl)-2-methoxybenzaldehyde (Intermediate 21). Int-19** (500 mg, 2.60 mmol), Cs_2_CO_3_ (1.69 g, 5.21 mmol), **Int-20** (562 mg, 3.12 mmol) and Pd(PPh_3_)_4_ (600 mg, 0.52 mmol) were dissolved in dioxane (5 ml) and H_2_O (1 ml), and the reaction was stirred at 100 °C for 2 hours under N_2._ The solvent was removed under reduced pressure and the residue was purified by flash chromatography on silica gel (PE/EA = 0 to 10 : 1) to afford **Int-21**: 600 mg, yield: 93%, (TLC: PE/EA = 10 : 1, Rf = 0.41). **LC-MS:** ([M+H]^+^ = 248). **^1^H NMR:** (CDCl_3_, 300 MHz) δ 8.49 (dd, *J* = 7.8, 1.8 Hz, 1H), 7.86 (dd, *J* = 7.6, 1.8 Hz, 1H), 7.35-7.31 (m, 1H), 3.95 (s, 3H), 2.43 (s, 3H), 2.41 (s, 3H).

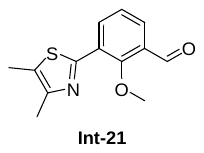

**Synthesis of 3-(4,5-dimethylthiazol-2-yl)-2-hydroxybenzaldehyde (Intermediate 22).** BBr_3_ in DCM (5ml) was added to **Int-21** (500 mg, 2.02 mmol) in DCM, and the reaction was stirred at room temperature for 3 hours, then sodium bicarbonate solution was added to pH 8 and extracted with DCM (100 mL × 3). The combined organic extracts were washed with brine (30 mL), dried over Na_2_SO_4_, filtered and the solvent was removed under reduced pressure. The residue was purified by flash chromatography on silica gel (DCM/MeOH = 0 to 10 : 1) to afford **Int-22**: 370 mg, yield: 79%, (TLC: DCM/MeOH = 10 : 1, Rf = 0.45). **LC-MS:** ([M+H]^+^ = 234).

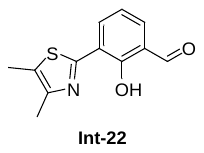

**Synthesis of 5-amino-1H-indazole-3-carboxylic acid (Intermediate 23).** **Int-23a** (200 mg, 0.97 mmol) was dissolved in MeOH (2 ml), Pd/C (20 mg) was added, and the reaction was stirred at 25 ^o^C for 12 hours under H_2_. The mixture was filtered, the solvent was removed under reduced pressure, and the residue was purified by flash chromatography on silica gel (DCM/MeOH = 0 to 10 : 1) to afford **Int-23**: 67 mg, yield: 39%, (TLC: DCM/MeOH = 10 : 1, Rf = 0.43). **LC-MS:** ([M+H]^+^ = 178).

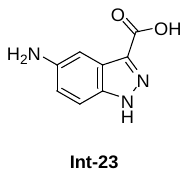

**Synthesis of 5-((3-(4,5-dimethylthiazol-2-yl)-2-hydroxybenzyl)amino)-1H-indazole-3-carboxylic acid (Compound 3). Int-22** (370 mg, 1.59 mmol) was dissolved in DMSO (5 ml), and **Int-23** (281 mg, 1.59 mmol), HOAc (96 mg, 1.59 mmol) and NaCNBH_3_ (400 mg, 6.36 mmol) were added, and the reaction was stirred at room temperature for 1 hour under N_2_. The residue was purified by flash chromatography on silica gel (DCM/MeOH = 0 to 10 : 1) to afford **Compound 3**: 30 mg, yield: 5%, (TLC: DCM/MeOH = 40 : 1, Rf = 0.36). **LC-MS:** ([M+H]^+^ = 395). **HPLC:** (purity: 92.71%). **^1^H NMR:** (CD_3_OD, 300 MHz) δ 13.35 (s, br, 1H), 12.63-12.41 (m, 2H), 7.53-7.48 (m, 1H), 7.40-7.32 (m, 2H), 7.08-6.93 (m, 2H), 6.87 (t, *J* = 7.7 Hz, 1H), 6.19 (s, 1H), 4.35 (s, 2H), 2.41 (s, 3H), 2.36 (s, 3H).

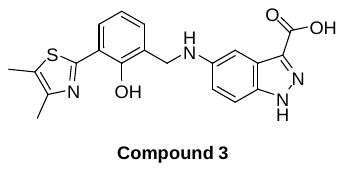

1.5. ***In vitro* characterisation of compounds 1 – 4.**
     1.5.1. **Surface plasmon resonance (SPR) binding assay.**

Surface plasmon resonance was performed as previously described^36,37^. Briefly, His-HAO1 (30 μg/mL) was immobilised on a Ni-NTA chip to a density of 5000 RU and equilibrated into the running buffer (20 mM HEPES, pH 7.5, 0.05% TWEEN20, 200 mM NaCl, 0.2% DMSO). A two-fold serial dilution (8 concentrations) was prepared in the above buffer for each analyte (0 – 2 μM for compounds **1**-**3** and 0 – 200 μM for compound **4**) and the subsequent solutions were passed over the chip at a flow rate of 30 μL/min. Data were plotted using GraphPad Prism software with curve fitting performed using a nonlinear least-squared regression fit to the Binding – Saturation: One Site – Total equation. Binding of each compound was tested in three independent experiments.

1.5.2. **Amplex Red activity assay.**

Inhibition of HAO1 was measured using an Amplex Red-based fluorescence assay to monitor hydrogen peroxide formation by HAO1 activity, as previously described^36,37^. All solutions were prepared in an assay buffer of 50 mM sodium phosphate, pH 7.4, 100 mM potassium chloride, 20 mM magnesium chloride and 0.01% Triton-X. Briefly, 30 nM HAO1 was incubated with varying compound concentrations (12 concentrations, 0 – 0.5 μM for compounds **1**-**3** and 0 – 500 μM for compound **4**) for five minutes before starting the reaction by addition of 30 μM glycolate and the Amplex Red assay reagent (100 μM Amplex Red dye (Cambridge Bioscience, catalogue no. CAY10010469), 0.2 U/mL horseradish peroxidase (Apollo Scientific, catalogue no. BITP1327), transferring into 384-well black assay plates (Greiner-One) and reading fluorescence (excitation 544 nm, emission 590 nm) on a POLARstar OMEGA plate reader (BMG Labtech). Inhibition of three different batches of recombinant His-HAO1 was measured for each compound with technical triplicates included for each batch. Data were plotted using GraphPad Prism software with curve fitting performed using a nonlinear least-squares regression fit to the log (inhibitor) vs response (three parameters) equation. Data plotted were mean values of the technical triplicates ± standard deviation.

1.5.3. **Thermal stabilisation by DSF.**

Differential scanning fluorimetry (DSF) was performed as previously described^36^ using a QuantStudio 5 RT-PCR machine (Applied Biosystems) with excitation and emission of 470 nm and 520 nm, respectively. All solutions were prepared in 50 mM HEPES, pH 7.5, 500 mM sodium chloride, 5% glycerol and 0.5 mM TCEP buffer (recombinant protein purification buffer). Briefly, 0.2 mg/mL recombinant His-HAO1 was added to varying compound concentrations (8 concentrations, 0 – 5 μM for compounds **1** and **2**, 0 – 100 μM for compound **3**, and 0 – 5 mM for compound **4**) before addition of 1:5000 SYPRO Orange dye (Invitrogen; catalogue no. 11510746). Samples were heated from 25 to 99 °C with a ramp rate of 0.05 °C/s, measuring fluorescence with every 1 °C increase. Multicomponent data were exported, transformed in excel to allow calculation of melting temperatures (T_m_) using the Boltzmann method, and plotted using GraphPad Prism software with curve fitting performed using a nonlinear least-squares regression fit to the log (inhibitor) vs response (three parameters) equation. Stabilisation of three different batches of recombinant His-HAO1 was measured for each compound with technical triplicates included for each batch. Data plotted were mean values of the technical triplicates ± standard deviation.

2. **Supplemental Figures and Tables**

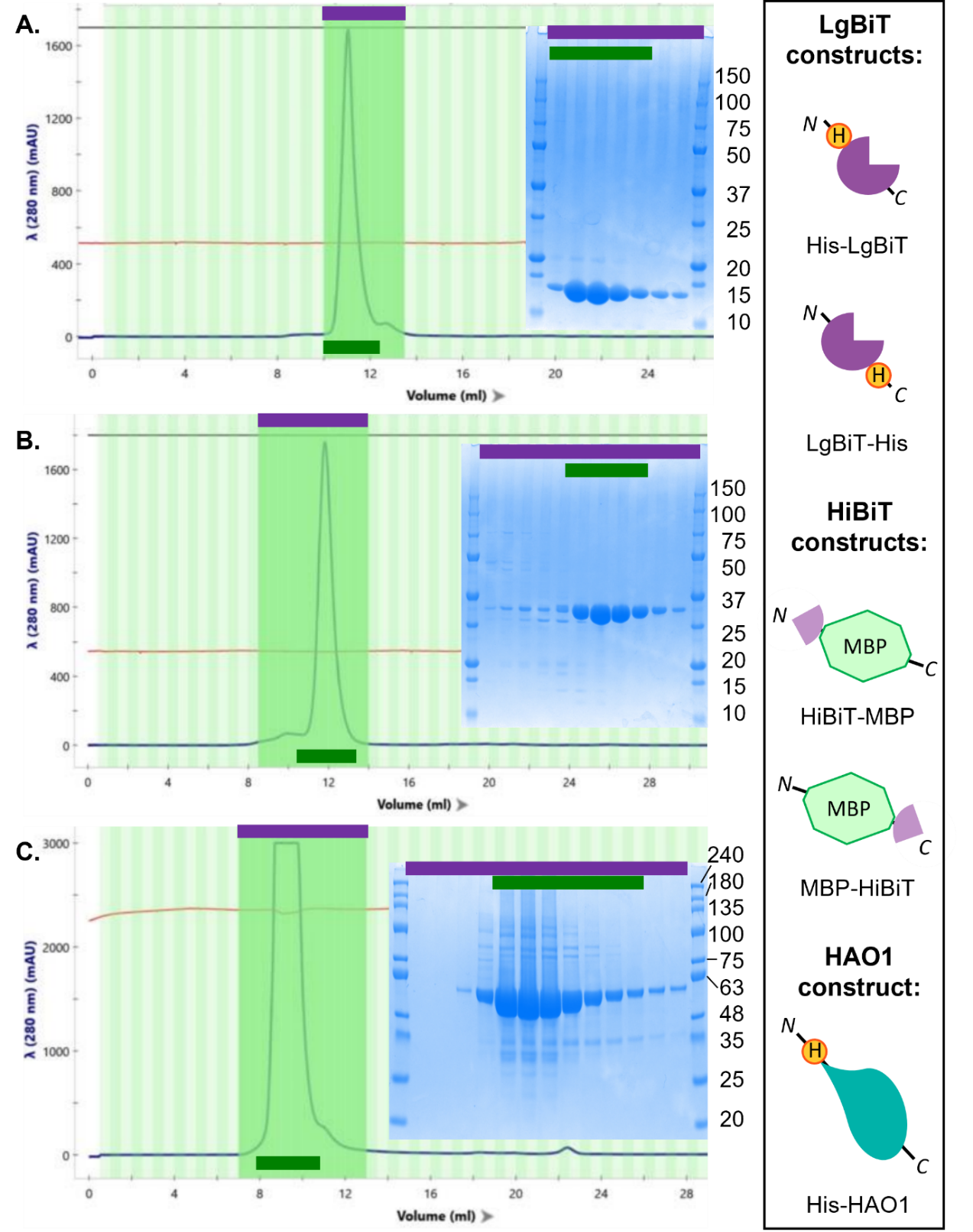

**Figure S1. Purification of recombinant protein SplitLuc components and controls.** Each panel shows the chromatogram trace (left) and SDS-PAGE analysis (right) for size exclusion chromatography performed on an S75 Increase column for His-LgBiT (A), HiBiT-MBP (B) and His-HAO1 (C) proteins. Chromatogram fractions that were analysed by SDS-PAGE are identified as a purple line on the chromatogram traces. Chromatogram fractions that were pooled to generate the final protein product are identified as a green line on both the chromatogram trace and the SDS-PAGE image. *Inset: Schematic illustrating the recombinant proteins, and the position of their incorporated tags, generated in this work.*

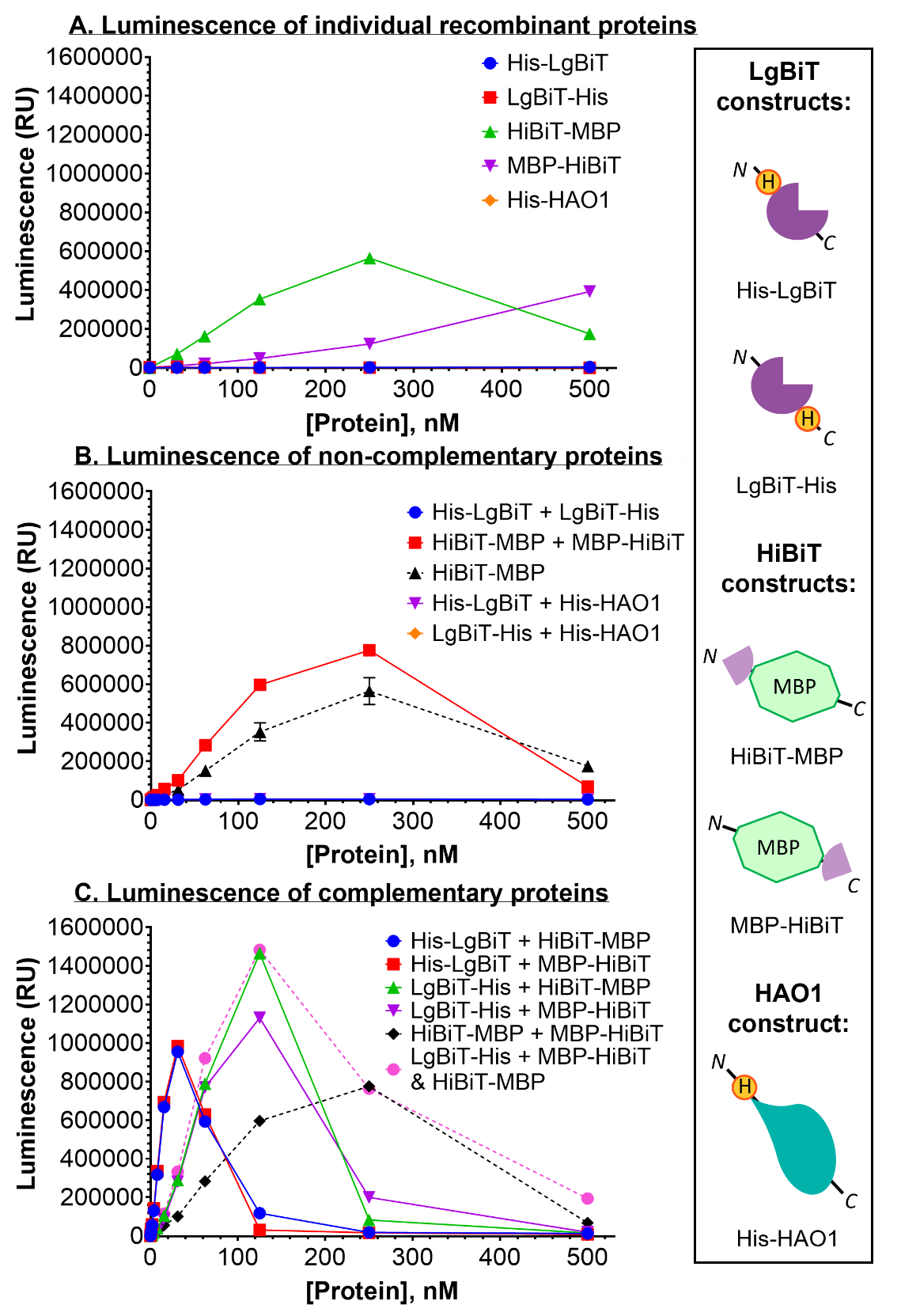

**Figure S2. Recombinant HiBiT-tagged proteins, but not His-HAO1, complement His-tagged LgBiT.** Data are represented as mean ±SD of technical triplicates. A. Luminescence of individual recombinant proteins as purified. B. Luminescence of each LgBiT protein mixed with either another LgBiT protein or His-HAO1 and of mixed HiBiT-tagged MBP proteins. Signal from HiBiT-MBP alone (panel A, green) is included as a dashed green line for reference. C. Luminescence of each LgBiT protein mixed with each HiBiT-tagged MBP protein. Signal from mixed HiBiT-tagged proteins (panel B, red) is included as a dashed red line for reference. Addition of luminescence values from LgBiT-His + MBP-HiBiT and HiBiT-MBP alone samples is also shown for reference (pink dashed line) and are not experimentally determined values. *Inset: Schematic illustrating recombinant proteins, and the position of their incorporated tags, characterised in this work.*

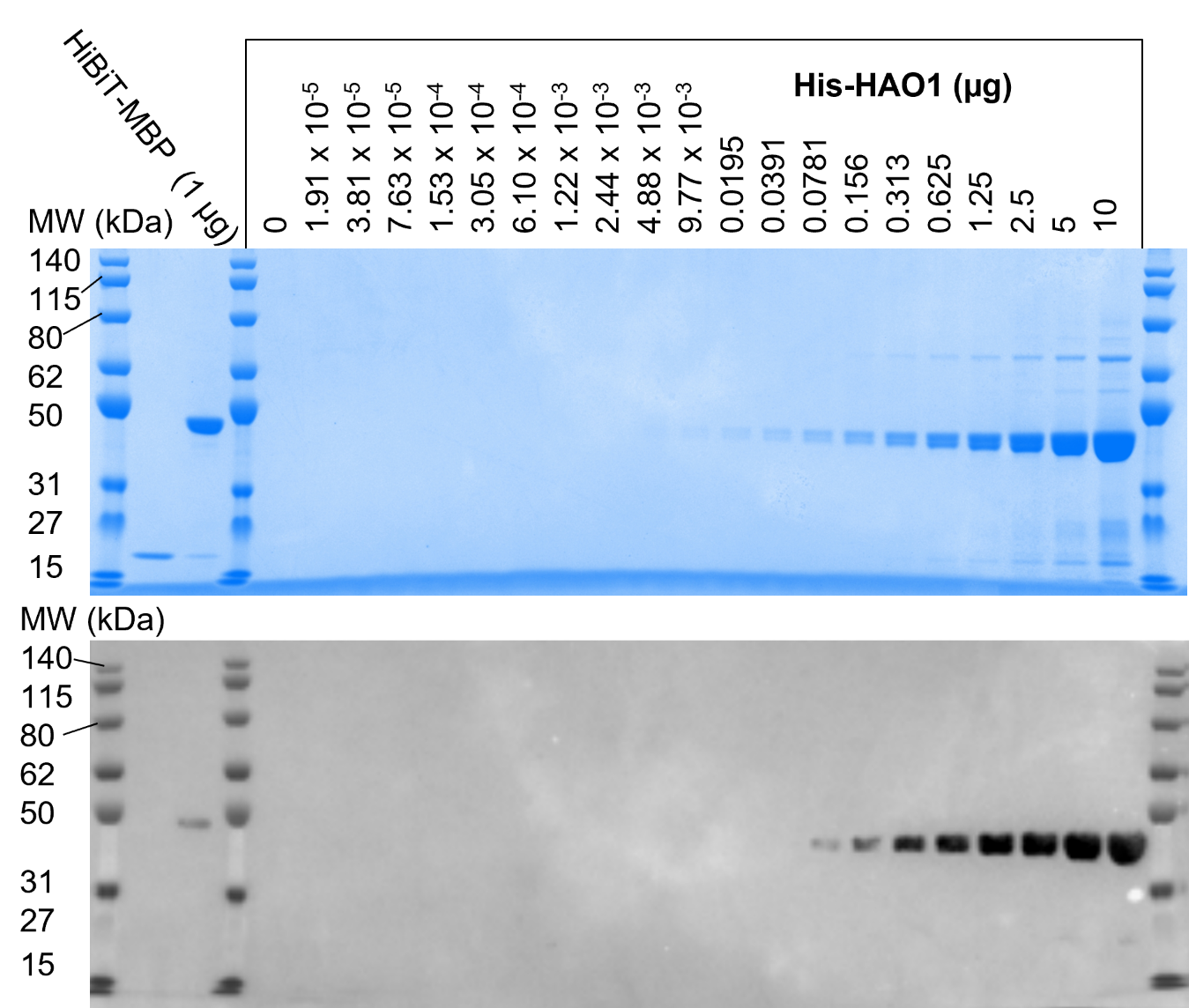
**Figure S3 Detection of recombinant His-HAO1, separated using SDS-PAGE, by Coomassie staining and Western Blot.** SDS-PAGE of two-fold serial dilution of recombinant His-HAO1 protein, prepared in SDS-PAGE loading dye, and HiBiT-MBP control, at 1 ug final amount (10 µL of 1 mg/mL stock). HAO1 protein was visualised by either addition of Coomassie gel stain and destaining with water (top) or transfer to a PVDF membrane, Western Blotting with anti-HAO1 primary antibody followed by anti-mouse IgG HRP-conjugated secondary antibody before addition of chemiluminescent HRP substrate (bottom), prior to imaging.

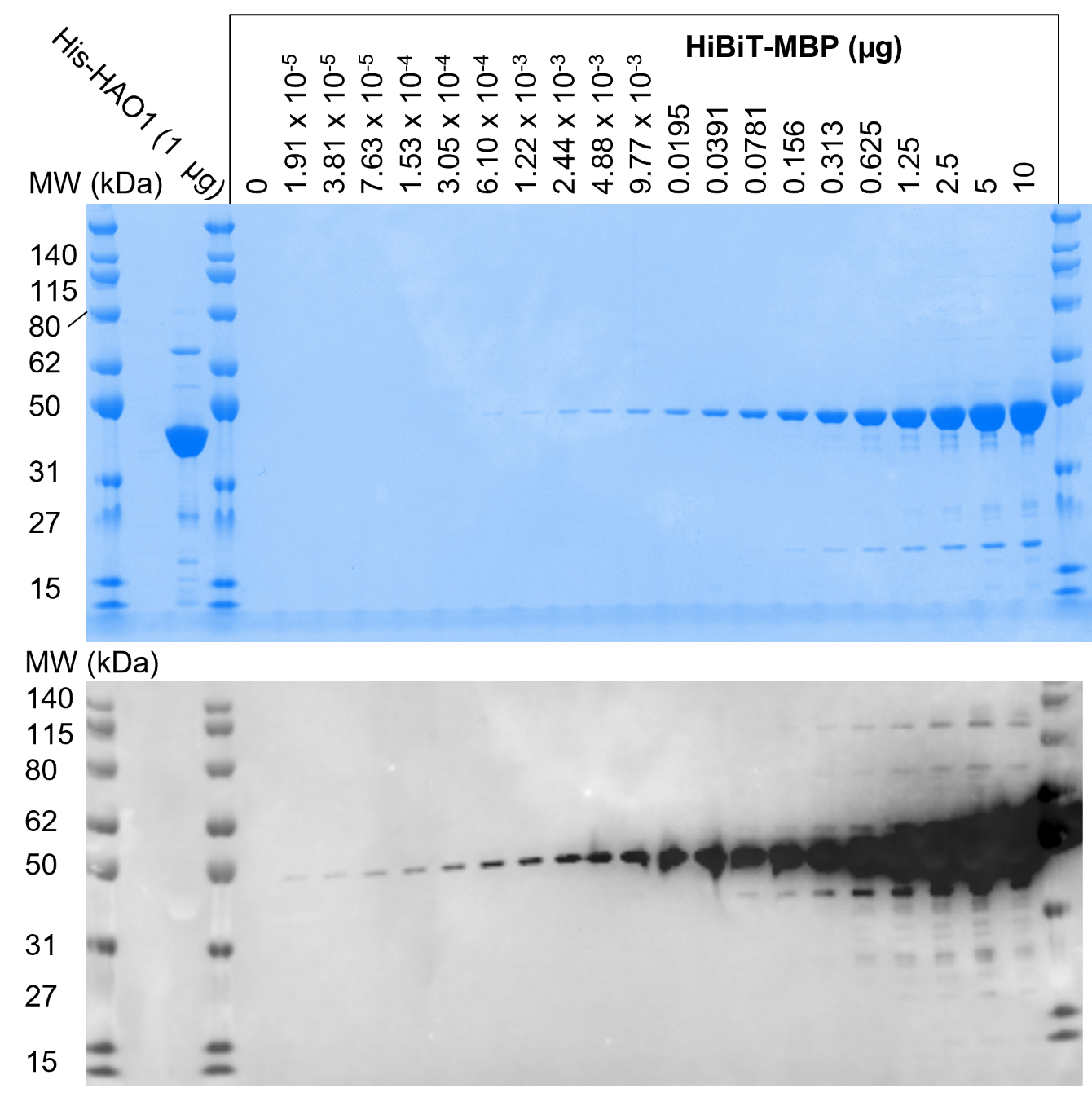

**Figure S4 Detection of recombinant HiBiT-MBP, separated using SDS-PAGE, by Coomassie staining and Western Blot.** SDS-PAGE of two-fold serial dilution of recombinant HiBiT-MBP protein, prepared in SDS-PAGE loading dye, and His-HAO1 control, at 1 ug final amount (10 µL of 1 mg/mL stock). HiBiT-MBP protein was visualised by either addition of Coomassie gel stain and destaining with water (top) or transfer to a PVDF membrane, Western Blotting with anti-HiBiT primary antibody followed by anti-mouse IgG HRP-conjugated secondary antibody before addition of chemiluminescent HRP substrate (bottom), prior to imaging.

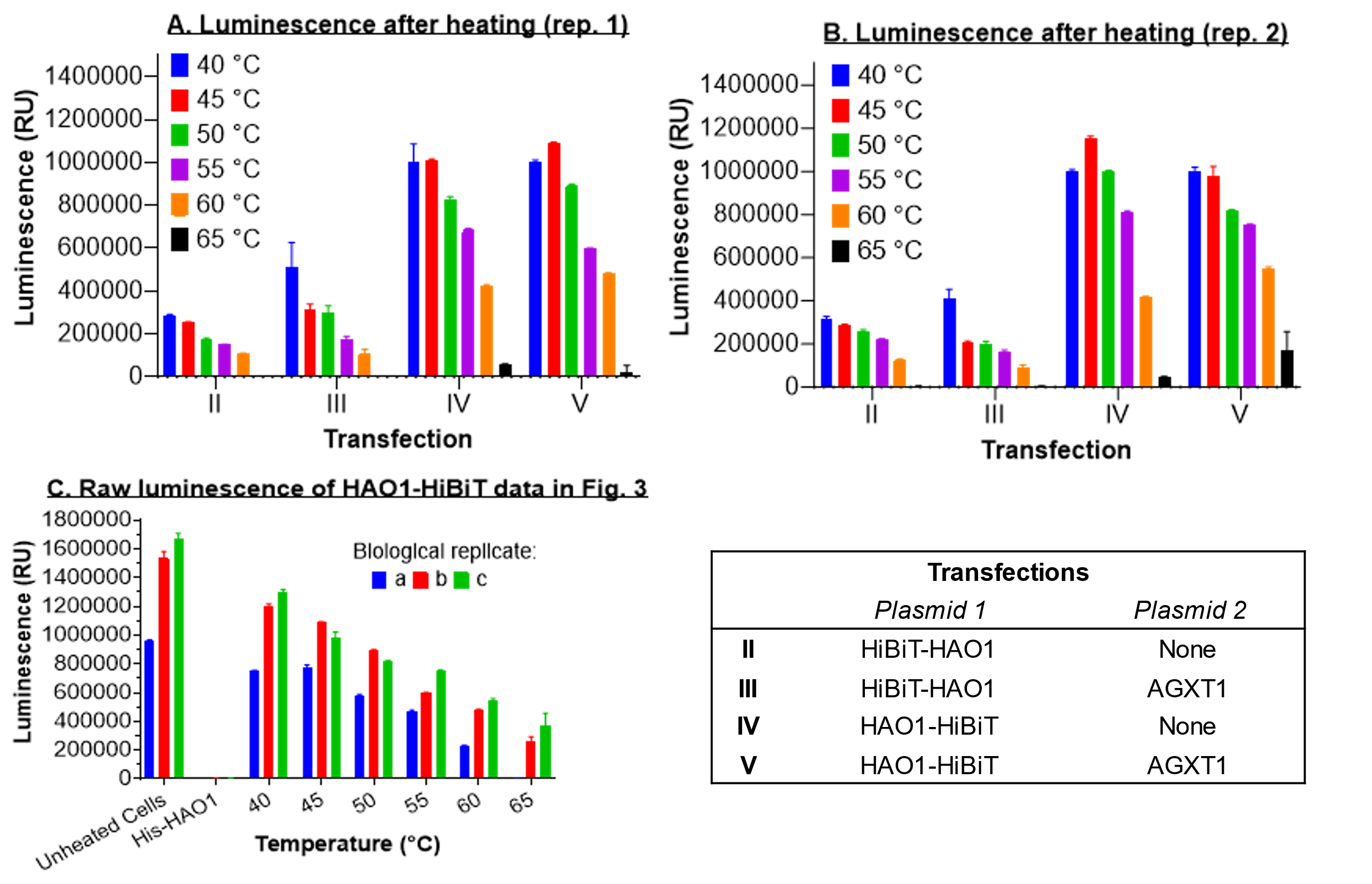

**Figure S5 Luminescence detection of HiBiT-tagged HAO1 from different transfections after heating between 40 and 65 °C.** Detection of soluble levels of HAO1-HiBiT protein, after heating at 5 °C intervals, by luminescence signal after addition of CETSA detection reagent (LgBiT protein and furimazine substrate). Data are represented as mean ± SD for technical triplicates. Panels A and B show testing of different transfections, each panel represents a different biological replicate. Panel C shows results for three different biological replicates of transfection III. Figure 3 in the main text represents this data after transformation to % Luminescence relative to the unheated control. *Inset: Definition of transfection mixes II*–*V.*

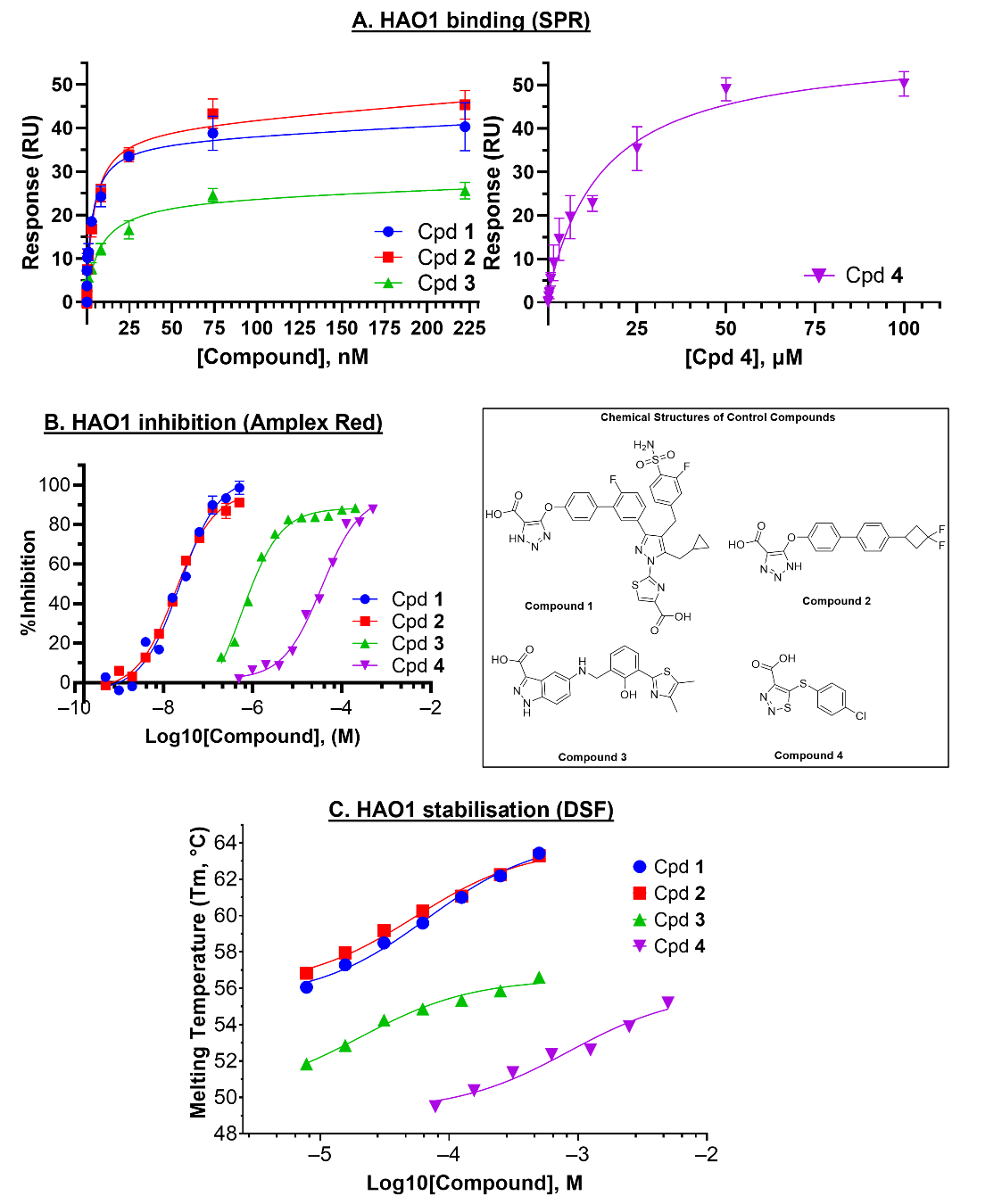

**Figure S6: *In vitro* inhibition, binding and stabilisation of recombinant His-HAO1 by control compounds.** Data are represented as mean ±SD of technical triplicates. A. Binding of HAO1 by compounds **1**–**4** measured using surface plasmon resonance. B. Inhibition of HAO1 by compounds **1**–**4** measured using Amplex Red activity assay. C. Thermal stabilisation of HAO1 by compounds **1**–**4** measured using differential scanning fluorimetry. *Inset: Chemical structures of tested compounds.*

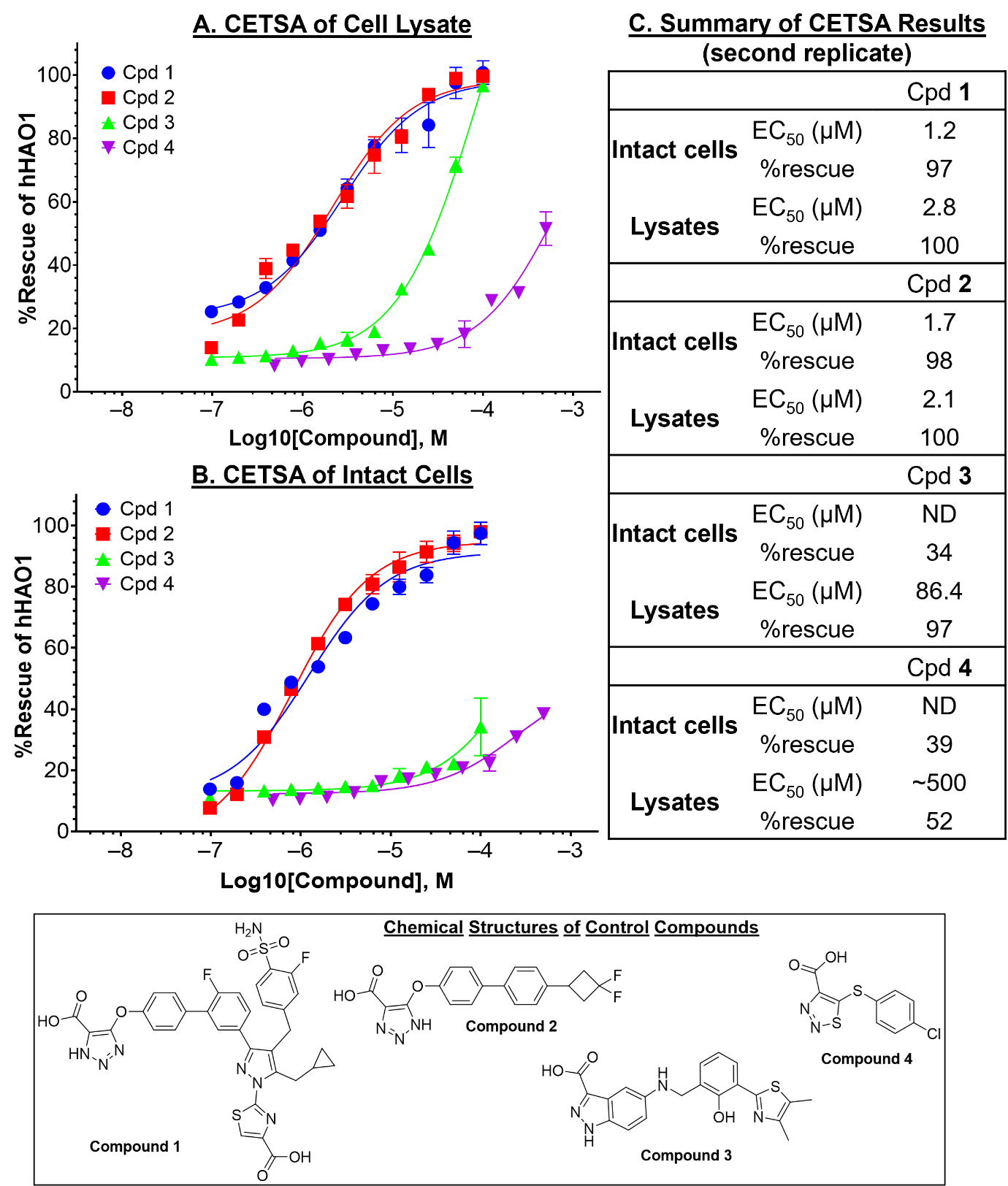

**Figure S7: Biological replicates of concentration-response testing of compounds 1–4 in SplitLuc CETSA.** Luminescence detection of soluble HAO1-HiBiT after preincubation with increasing compound concentrations and heating to 55 °C, measured in either intact cells (A) or cell lysates (B). Data are represented as mean ±SD of technical triplicates. C. Summary of compound parameters determined in CETSA experiments. *Inset: Chemical structures of tested compounds.*

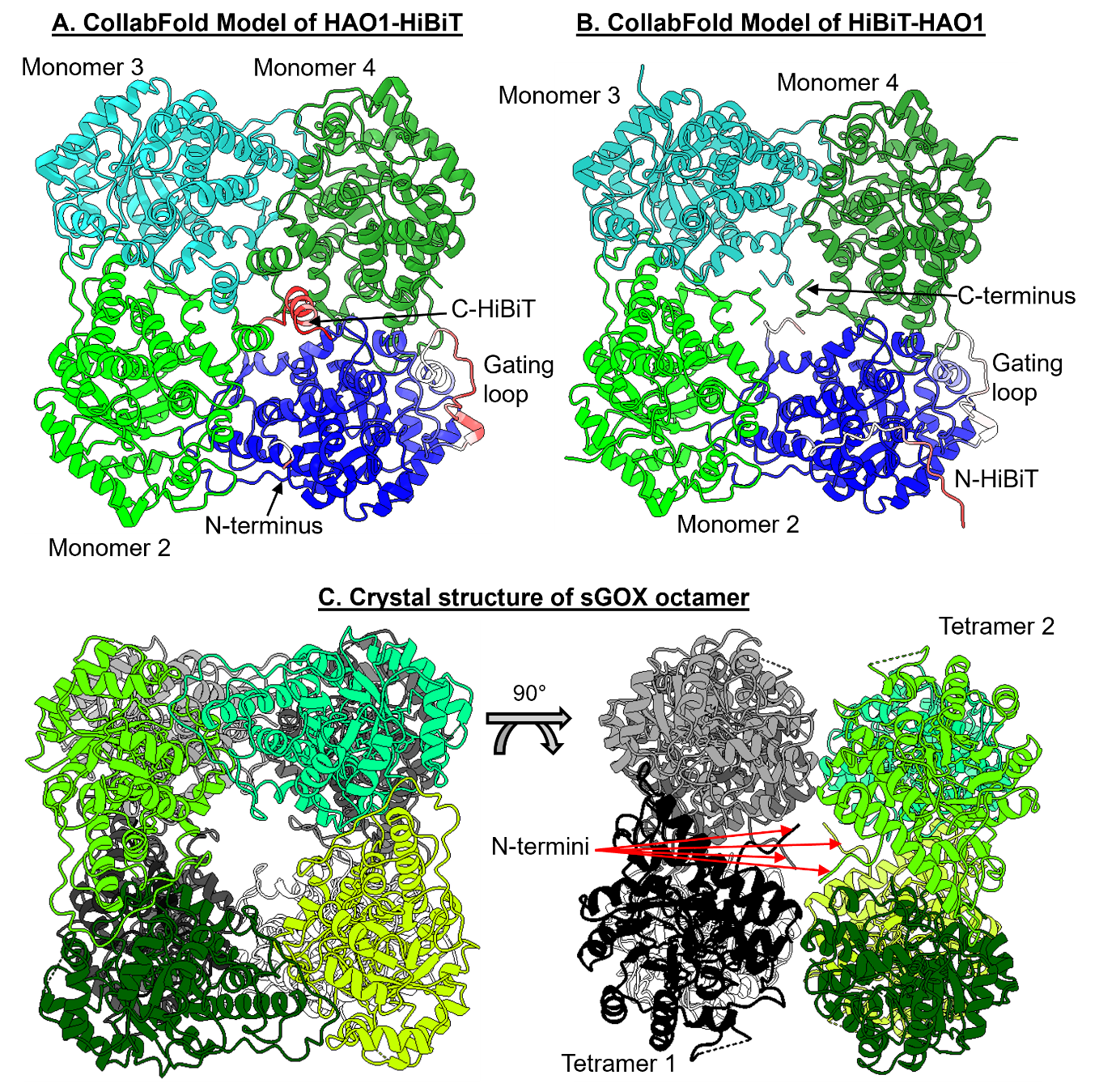

**Figure S8. Structural models of possible oligomers of HiBiT-tagged HAO1 and the spinach homologue of HAO1, sGOX.** Structural models of HAO1-HiBiT (A) and HiBiT-HAO1 (B) tetramers, generated through the ColabFold functionality of Chimera-X. In each panel, one monomer is shaded according to the per-residue measure of local confidence (predicted local distance difference test, pLDDT) provided by the AlphaFold algorithm, from high confidence (blue) to low confidence (red). Note that the two areas of low confidence were the HiBiT tag and the flexible, and often disordered, gating loop. The other three monomers are shown as light green, light blue and dark green ribbons. C. Octameric arrangement of spinach glycolate oxidase, sGOX, within the published crystal structure (PDB 1gox). Monomers of one tetramer are shown as ribbons of varying shades of green and monomers of the second tetramer are shown as ribbons of varying shades of grey. The image to the left is shown from the same orientation as the HiBiT-tagged HAO1 tetramers in panels A and B. Rotation of the left hand structure by 90 ° allows visualisation of the interaction between the N-termini of the two different tetramers, as shown in the right hand structure.

**Table S1: Vector and sequence details of constructs used in this study.** Related to Figure 1.

| **Construct** | **Vector** | **Protein sequence** |
| --- | --- | --- |
| *E. coli expression plasmids* | | |
| His-HAO1 | pNIC28-Bsa4 | MHHHHHHSSGVDLGTENLYFQSMLPRLICINDYEQHAKSVLPKSIYDYYRSGANDEETLADNIAAFSRWKLYPRMLRNVAETDLSTSVLGQRVSMPICVGATAMQRMAHVDGELATVRACQSLGTGMMLSSWATSSIEEVAEAGPEALRWLQLYIYKDREVTKKLVRQAEKMGYKAIFVTVDTPYLGNRLDDVRNRFKLPPQLRMKNFETSTLSFSPEENFGDDSGLAAYVAKAIDPSISWEDIKWLRRLTSLPIVAKGILRGDDAREAVKHGLNGILVSNHGARQLDGVPATIDVLPEIVEAVEGKVEVFLDGGVRKGTDVLKALALGAKAVFVGRPIVWGLAFQGEKGVQDVLEILKEEFRLAMALSGCQNVKVIDKTLVRKNPLAVSKI |
| His-LgBiT | pNIC28-Bsa4 | MHHHHHHSSGVDLGTENLYFQSMVFTLEDFVGDWEQTAAYNLDQVLEQGGVSSLLQNLAVSVTPIQRIVRSGENALKIDIHVIIPYEGLSADQMAQIEEVFKVVYPVDDHHFKVILPYGTLVIDGVTPNMLNYFGRPYEGIAVFDGKKITVTGTLWNGNKIIDERLITPDGSMLFRVTINS |
| LgBiT-His | pNIC28-CTH0 | MVFTLEDFVGDWEQTAAYNLDQVLEQGGVSSLLQNLAVSVTPIQRIVRSGENALKIDIHVIIPYEGLSADQMAQIEEVFKVVYPVDDHHFKVILPYGTLVIDGVTPNMLNYFGRPYEGIAVFDGKKITVTGTLWNGNKIIDERLITPDGSMLFRVTINSAENLYFQSHHHHHH |
| HiBiT-MBP | pNIC28-CTH0 | MGSVSGWRLFKKISGSGGSKIEEGKLVIWINGDKGYNGLAEVGKKFEKDTGIKVTVEHPDKLEEKFPQVAATGDGPDIIFWAHDRFGGYAQSGLLAEITPDKAFQDKLYPFTWDAVRYNGKLIAYPIAVEALSLIYNKDLLPNPPKTWEEIPALDKELKAKGKSALMFNLQEPYFTWPLIAADGGYAFKYENGKYDIKDVGVDNAGAKAGLTFLVDLIKNKHMNADTDYSIAEAAFNKGETAMTINGPWAWSNIDTSKVNYGVTVLPTFKGQPSKPFVGVLSAGINAASPNKELAKEFLENYLLTDEGLEAVNKDKPLGAVALKSYEEELAKDPRIAATMENAQKGEIMPNIPQMSAFWYAVRTAVINAASGRQTVDEALKDAQTSSGHHHHHHHHHH |
| MBP-HiBiT | pNIC28-CTH0 | MHHHHHHHHHHGGSKIEEGKLVIWINGDKGYNGLAEVGKKFEKDTGIKVTVEHPDKLEEKFPQVAATGDGPDIIFWAHDRFGGYAQSGLLAEITPDKAFQDKLYPFTWDAVRYNGKLIAYPIAVEALSLIYNKDLLPNPPKTWEEIPALDKELKAKGKSALMFNLQEPYFTWPLIAADGGYAFKYENGKYDIKDVGVDNAGAKAGLTFLVDLIKNKHMNADTDYSIAEAAFNKGETAMTINGPWAWSNIDTSKVNYGVTVLPTFKGQPSKPFVGVLSAGINAASPNKELAKEFLENYLLTDEGLEAVNKDKPLGAVALKSYEEELAKDPRIAATMENAQKGEIMPNIPQMSAFWYAVRTAVINAASGRQTVDEALKDAQTSSGVDLGTENLYFQ\SGSVSGWRLFKKISGS |

**Table S1 (continued): Vector and sequence details of constructs used in this study.** Related to Figure 1.

| **Construct** | **Vector** | **Protein sequence** |
| --- | --- | --- |
| *Mammalian expression plasmids* | | |
| HiBiT-HAO1 | pHTBV1.1 | MGSVSGWRLFKKISGSLPRLICINDYEQHAKSVLPKSIYDYYRSGANDEETLADNIAAFSRWKLYPRMLRNVAETDLSTSVLGQRVSMPICVGATAMQRMAHVDGELATVRACQSLGTGMMLSSWATSSIEEVAEAGPEALRWLQLYIYKDREVTKKLVRQAEKMGYKAIFVTVDTPYLGNRLDDVRNRFKLPPQLRMKNFETSTLSFSPEENFGDDSGLAAYVAKAIDPSISWEDIKWLRRLTSLPIVAKGILRGDDAREAVKHGLNGILVSNHGARQLDGVPATIDVLPEIVEAVEGKVEVFLDGGVRKGTDVLKALALGAKAVFVGRPIVWGLAFQGEKGVQDVLEILKEEFRLAMALSGCQNVKVIDKTLVRKNPLAVS |
| HAO1-HiBiT | pHTBV1.1 | MLPRLICINDYEQHAKSVLPKSIYDYYRSGANDEETLADNIAAFSRWKLYPRMLRNVAETDLSTSVLGQRVSMPICVGATAMQRMAHVDGELATVRACQSLGTGMMLSSWATSSIEEVAEAGPEALRWLQLYIYKDREVTKKLVRQAEKMGYKAIFVTVDTPYLGNRLDDVRNRFKLPPQLRMKNFETSTLSFSPEENFGDDSGLAAYVAKAIDPSISWEDIKWLRRLTSLPIVAKGILRGDDAREAVKHGLNGILVSNHGARQLDGVPATIDVLPEIVEAVEGKVEVFLDGGVRKGTDVLKALALGAKAVFVGRPIVWGLAFQGEKGVQDVLEILKEEFRLAMALSGCQNVKVIDKTLVRKNPLAVSGSVSGWRLFKKISGS |
| AGXT1 | pHTBV1.1-CT10H-SIII-LIC | MASHKLLVTPPKALLKPLSIPNQLLLGPGPSNLPPRIMAAGGLQMIGSMSKDMYQIMDEIKEGIQYVFQTRNPLTLVISGSGHCALEAALVNVLEPGDSFLVGANGIWGQRAVDIGERIGARVHPMTKDPGGHYTLQEVEEGLAQHKPVLLFLTHGESSTGVLQPLDGFGELCHRYKCLLLVDSVASLGGTPLYMDRQGIDILYSGSQKALNAPPGTSLISFSDKAKKKMYSRKTKPFSFYLDIKWLANFWGCDDQPRMYHHTIPVISLYSLRESLALIAEQGLENSWRQHREAAAYLHGRLQALGLQLFVKDPALRLPTVTTVAVPAGYDWRDIVSYVIDHFDIEIMGGLGPSTGKVLRIGLLGCNATRENVDRVTEALRAALQHCPKKKLAENLYFQSHHHHHHHHHHGSAWSHPQFEKGGGSGGGSGGSAWSHPQFEK |

**Table S2: Published characterisation data for compounds 1 – 4 in recombinant protein and cellular assays**. ^a^First IC_50_ was measured in wild type mouse hepatocytes; second IC_50_ was measured in AGXT knockdown mouse hepatocytes. ^b^First IC_50_ was measured in mouse AGXT^-/-^ hepatocytes after 24 hours; second IC_50_ is measured after 48 hours. ^c^Rescue of glycolate-induced toxicity in hHAO1 transfected CHO cells. ^d^Reduction in urinary oxalate excretion after 24 hours, measured in mouse AGXT^-/-^ hepatocytes. Related to Figure 4.

|  | **Recombinant Protein Assays** | | **Cellular Assays** | **Reference(s)** |
| --- | --- | --- | --- | --- |
|  | **Potency (IC_50_)** | **Affinity** |  |  |
| **1** | 4.7 nM | ND | IC_50_ 136 nM/ 88 nM^a^ | [S1] |
| **2** | 15.4 nM | K_d_ 6.31 nM | IC_50_ 24.2 nM/ 42.9 nM^b^ | [S2, S3] |
| **3** | 25 nM | ND | 76% rescue at 30 µM^c^ | [S4] |
| **4** | 22 µM | K_D_ 47.5 µM | EC_50_ 25.26 µM; max. 70% oxalate reduction^d^ | [S5, S6, S7] |

3. **Supplemental References**

[S1] Ding, J., Gumpena, R., Boily, M., Caron, A., Chong, O., Cox, J.H., Dumais, V., Gaudreault, S., Graff, A.H., King, A. et al. (2021). Dual Glycolate Oxidase/Lactate Dehydrogenase A Inhibitors for Primary Hyperoxaluria. ACS Med Chem Lett. 12 (7): 1116-1123. doi:10.1021/acsmedchemlett.1c00196.

[S2] Maag, H., Fernandes, M.X., Zamboni, R., Akbariromani, E., Beaulieu, M.A., Leblanc, Y., and Thakur, P. (2020). Triazole glycolate oxidase inhibitors. World Intellectual Property Organization, WO2020010309A1. https://patentscope.wipo.int/search/en/detail.jsf?docId=WO2020010309

[S3] Sinha, U., Rajasingham, T., Xavier Fernandez, M., Ji, A., Sukhun, R., Rao, S., Henry, C., D’souza, K., and Lafountaine, J. (2022). MO040 Development of BBP-711, A small molecule inhibitor of glycolate oxidase for primary hyperoxaluria type 1 and recurrent kidney stone formers. Nephrology Dialysis Transplantation *37*. https://doi.org/10.1093/ndt/gfac062.021.

[S4] Lee, E.C.Y., McRiner, A.J., Georgiadis, K.E., Liu, J., Wang, Z., Ferguson, A.D., Levin, B., von Rechenberg, M., Hupp, C.D., Monterio, M.I. et al. (2021). Discovery of Novel, Potent Inhibitors of Hydroxy Acid Oxidase 1 (HAO1) Using DNA-Encoded Chemical Library Screening. J Med Chem. 64 (10): 6730-6744. doi:10.1021/acs.jmedchem.0c02271.

[S5] Chen, Z., Vignaud, C., Jaafar, A., Levy, B., Gueritte, F., Guenard, D., Lederer, F. and Mathews, F.S. (2012). High resolution crystal structure of rat long chain hydroxy acid oxidase in complex with the inhibitor 4-carboxy-5-[(4-chlorophenyl)sulfanyl]-1, 2, 3-thiadiazole. Implications for inhibitor specificity and drug design. Biochimie. 94 (5): 1172-1179. doi:10.1016/j.biochi.2012.02.003.

[S6] Martin-Higueras, C., Luis-Limas, S., and Salido, E. (2016). Glycolate Oxidase Is a Safe and Efficient Target for Substrate Reduction Therapy in a Mouse Model of Primary Hyperoxaluria Type I. Mol Ther. 24 (4): 719-725. doi:10.1038/mt.2015.224.

[S7] Mackinnon, S.R., Bezerra, G.A., Krojer, T., Szommer, T., von Delft, F., Brennan, P.E., and Yue, W.W. (2022). Novel Starting Points for Human Glycolate Oxidase Inhibitors, Revealed by Crystallography-Based Fragment Screening. Front Chem *10*. https://doi.org/10.3389/fchem.2022.844598.
